# Large scale analysis of predicted protein structures links model features to *in vivo* behaviour

**DOI:** 10.1101/2024.04.10.588835

**Authors:** Michael J. Stam, Diego A. Oyarzún, Nadanai Laohakunakorn, Christopher W. Wood

## Abstract

Rapid advancements in protein structure prediction methods have ushered in a new era of abundant and accurate structural data, providing opportunities to analyse proteins at a scale that has not been possible before. Here we show that features derived solely from predicted structures can be used to understand *in vivo* protein behaviour using data-driven methods. We found that these features were predictive of *in vivo* protein production for a set of designed antibodies, enabling identification of high-quality designs. Following on from this result, we calculated these features for a diverse set of ≈500,000 predicted structures, and our analysis showed systematic variation between proteins from different organisms to such an extent that the tree of life could be recapitulated from these data. Given the high degree of functional constraint around the chemistry of proteins, this result is surprising, and could have important implications for the design and engineering of novel proteins.

## Introduction

The development of highly accurate protein structure prediction methods, such as AlphaFold (Jumper et al., 2021; Senior et al., 2020) and ESMFold (Lin et al., 2023) has resulted in unprecedented amounts of protein structural data becoming available to researchers. The AlphaFold DB (Varadi et al., 2022) has over 200 million predicted protein structures, covering the majority of the UniProt database (Consortium, 2023) and a huge variety of organisms. Complementary to the AlphaFold DB is the ESM Metagenomic Atlas (Lin et al., 2023) which used ESMFold to obtain predicted structures for 772 million metagenomic proteins, and the MIP database (Koehler Leman et al., 2023) which used Rosetta (Leman et al., 2020) and DMPFold (Greener et al., 2019) to predict the structures of 200,000 metagenomic proteins across the microbial tree of life. By leveraging these large structural data sets, researchers can now study the biological properties and functions of proteins at a scale that has never been possible before. Furthermore, the insight gained from exploring these data sets is important for understanding the role of proteins in nature and designing novel proteins to address challenges across various scientific areas.

Recent studies have used these large structural data sets for a broad range of applications including; uncovering new protein families and folds (Durairaj et al., 2023), identifying and prioritising drug targets (Ochoa et al., 2023), augmenting training data to learn inverse folding for protein sequence design (Hsu et al., 2022), and improving phylogenetic trees (Moi et al., 2023). These studies show a glimpse of the potential that these data sets have in the fields of structural biology, protein design, medicine, and evolutionary biology, and they demonstrate that we are at the beginning of a completely new field of science (Callaway, 2023).

Here, we performed a large-scale analysis of predicted protein structures to determine if physicochemical descriptors of structural models were predictive of *in vivo* behaviour. We first calculated a set of features using our structural model evaluation server DE-STRESS (Stam & Wood, 2021). This programme calculates roughly 70 structural features using a range of software including all-atom scoring functions, geometric measures of packing density and hydrogen bonding quality, aggregation propensity, isoelectric point and many more (Figure 1). These properties were determined for 192 designed single chain variable fragment (scFv) antibodies from the Fleishman lab (Baran et al., 2017). Unsupervised and supervised learning methods were used to demonstrate that these properties were predictive of protein production. Following on from this, we applied similar methods to 564,446 AlphaFold2 (AF2) models from 48 model organisms (Varadi et al., 2022), to gain an understanding of how these properties varied at scale. Finally, we found that there were systematic differences in the physicochemical properties between organisms, to such an extent that these properties could be used to reconstruct the tree of life.

**Figure 1.**
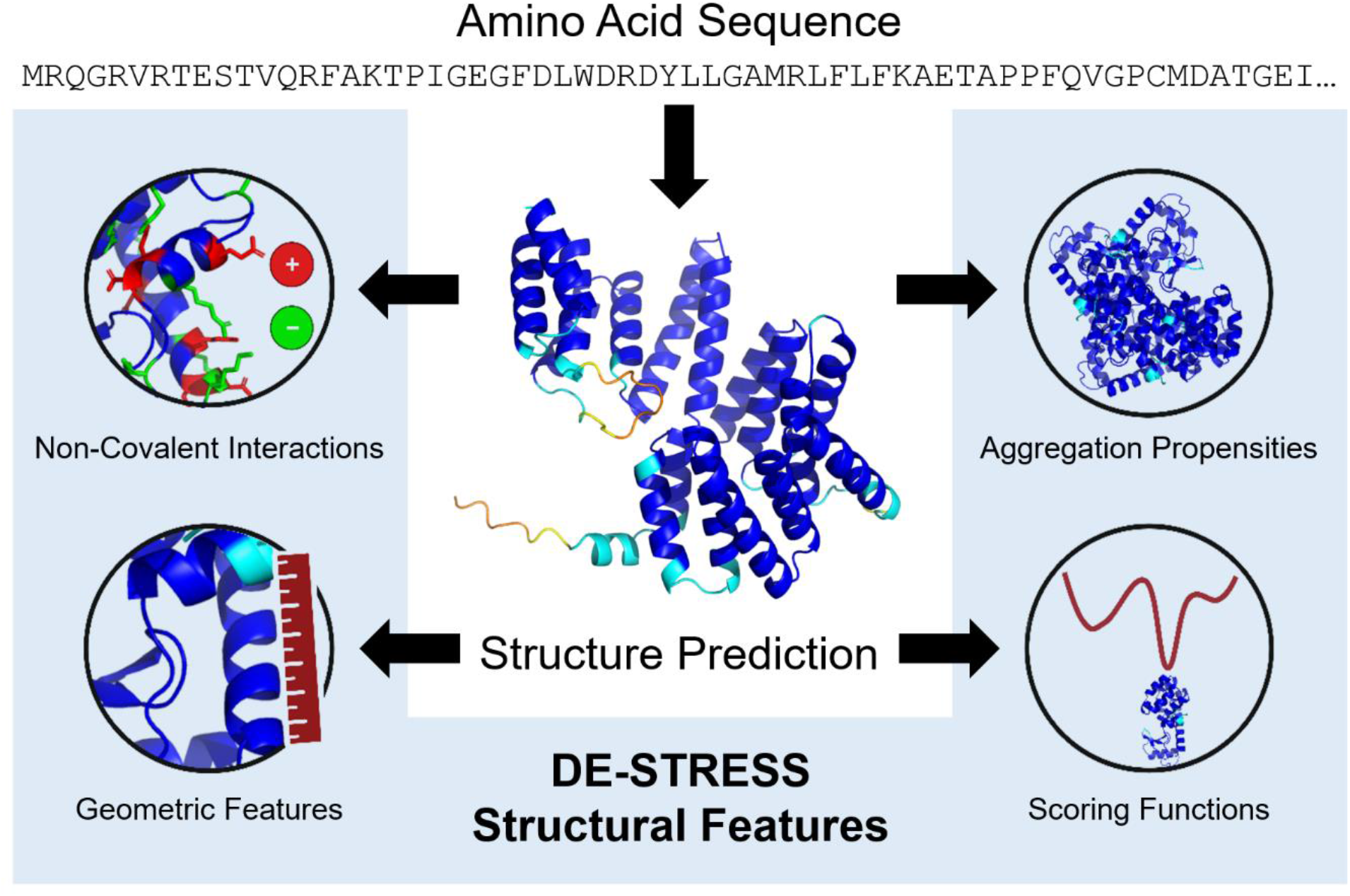
Overview of the computational pipeline detailing how AlphaFold and DE-STRESS are used to extract physicochemical properties from structural models of proteins.

## Results

### Physicochemical Properties Can Be Used to Predict Protein Production Levels

We analysed a set of designed single chain variable fragment (scFv) antibodies, which had been designed and experimentally characterised previously (Baran et al., 2017) (Figure 2A). This set of proteins was of interest as, while sequences were generated using the same basic design method, they showed vastly different levels of protein production experimentally. AlphaFold2 (Jumper et al., 2021) was used to generate structural models of these proteins, and DE-STRESS (Stam & Wood, 2021) was then used to calculate a set of physicochemical properties. A glossary of the different DE-STRESS metrics used in this work are detailed in supplementary tables S1 to S8. After generating this data set, we performed dimensionality reduction using principal component analysis (PCA) (Pearson, 1901). A plot of the cumulative variance explained by each principal component is shown in Figure S1 and the major contributors to the top two principal components (PC1 and PC2) are shown in Table S9. Plotting PC1 and PC2 against each other shows multiple clusters that separate out protein production into low, low/medium, and high-level clusters for the designed antibodies (Figure 2B). The structural predictions for unrelated experimentally determined scFvs, obtained from the Structural Antibody Database (SAbDab) (Dunbar et al., 2014; Schneider et al., 2022), cluster together with the highly produced designs, indicating that they share features that are related to this property. PC1 and PC2 explain over 50% of the variance in the data set, with aggregation propensity metrics (Kuriata et al., 2019) and lysine, glycine and aspartate proportions contributing the most to PC1, while glycine, glutamine and valine proportions and energy scores capturing hydrogen bonds between side chains and backbone (Alford et al., 2017), contributing the most to PC2 (Table S9). Figure S2 repeats the PCA analysis with the amino acid composition metrics excluded from the data set, and similar results are observed.

**Figure 2.**
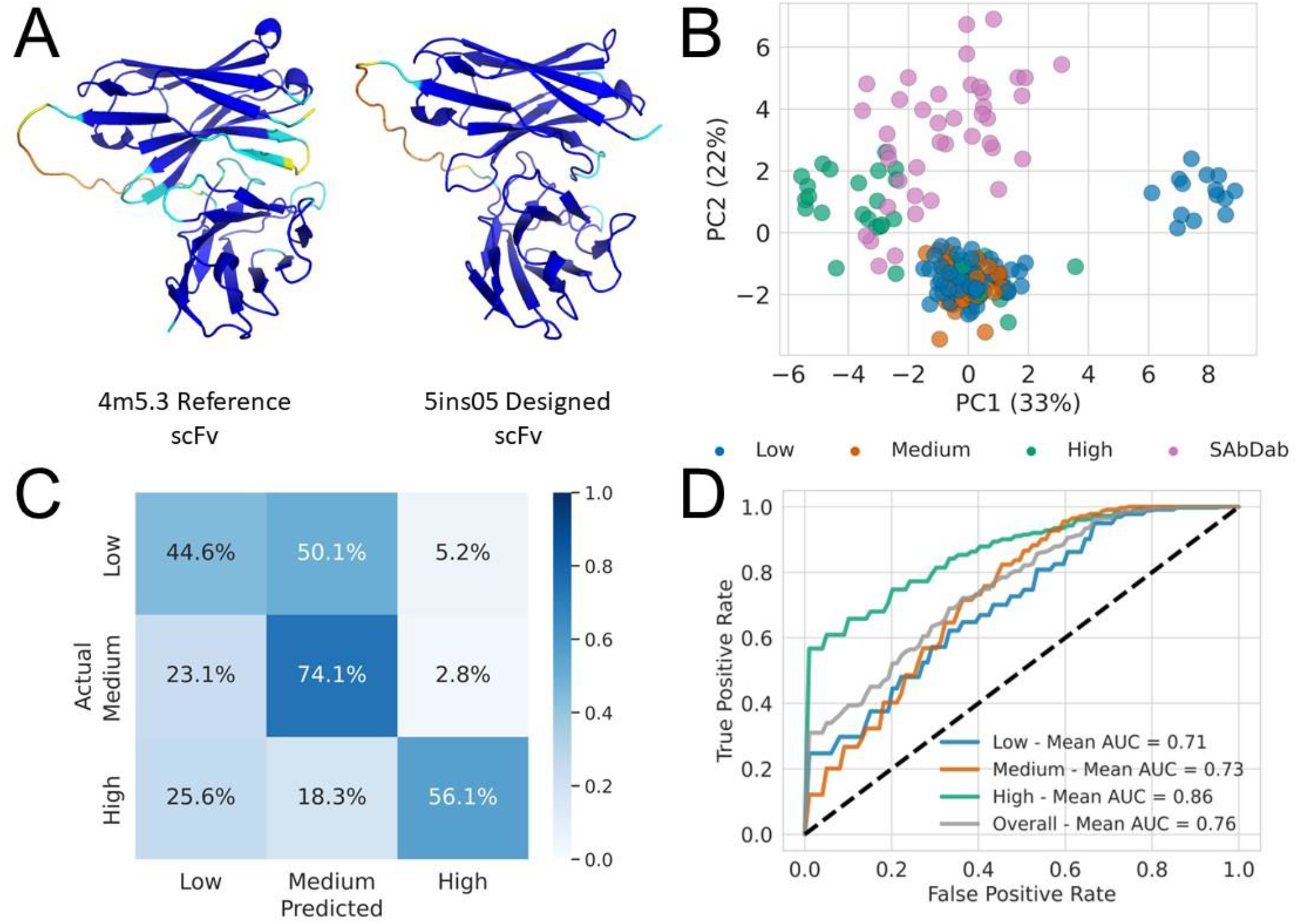
A) AlphaFold2 structural models of the reference scFv and a designed scFv from the Fleishman dataset. B) A plot of PC1 and PC2 for the scFv designs and 41 experimentally determined scFvs, along with the variance explained. C) A confusion matrix for the best classifier. D) ROC curves split out by protein production level, and an overall ROC curve, for the best classifier.

After exploring these data with PCA, we attempted to classify levels of protein production using simple Naive Bayes classifiers (Bishop, 2006) (figure 2C/D). The best classifier had mean ROC AUC of 0.76 across the repetitions of 5-fold cross validation. In addition to this, the rest of the classifiers fitted across the folds, still performed well with mean ROC AUCs ranging from 0.73 to 0.76 (Table S10). The top performing model correctly classifies 56% of high producing designs and incorrectly predicts the rest as low or medium producing. In contrast to this, very few of the low and medium producing designs are misclassified as high producing, a feature that would be useful when triaging protein designs to be characterised in the lab. Next, the performance of this classifier was evaluated on the test set, and it had weighted ROC AUC of 0.78, weighted precision of 0.68 and weighted recall of 0.60, showing that it generalises well to unseen data (Table S11). Features that were important for the prediction accuracy included certain amino acid composition metrics, hydrophobic fitness (E. S. Huang et al., 1995), aggregation propensity scores (Kuriata et al., 2019), and Rosetta scores capturing solvation energy (Alford et al., 2017) which are shown in tables S12 and S13 and ordered by feature importance, with the highest importance at the top and lowest at the bottom.

### Large-Scale Analysis of Physicochemical Properties Performed Across Half a Million Predicted Protein Structures

Following on from the analysis of the small antibody data set, we decided to apply these properties to a large data set of structural models from the AlphaFold DB, to gain greater insight into how the physicochemical properties varied between protein models. We calculated DE-STRESS metrics for 564,446 AF2 structural models, excluding amino acid composition metrics to be sure that any systematic variation was not the result of sequence bias, and PCA was performed on these features scaled using the minmax scaling method. PC1 and PC2 accounted for over 50% of the variance from the original data set (Figure 3A). One observation from this scatter plot is that there were clear differences in secondary structure across this space, with α-helix rich proteins to the top of the plot, β-rich proteins to the bottom right and disordered proteins to the bottom left, while proteins in the middle had a mix of secondary structure types (Figure 3A/B). Some of the main contributors to PC2 were consistent with secondary structure, including long and short-range hydrogen bond energy values and Rosetta energy terms capturing rotamer preferences (Table 1). Isoelectric point also varied across the space (Figure 3C), along with aggregation propensity metrics, and long and short-range hydrogen bond energies which are shown to be the main contributors to PC1 (Table 1). Cumulative histograms for secondary structure, isoelectric point, packing density and aggregation propensity across PC1 and PC2 demonstrate these relationships clearly (Figure S4). This analysis was repeated using the robust scaling method and on the Protein Data Bank (PDB) (Berman et al., 2000), rather than predicted structures (Figures S5-10). While the results were generally consistent, when the robust scaling method was used, packing density and aggregation propensity scores were the major factors that varied across these spaces and there was no clear relationship with isoelectric point (Tables S14-16).

**Table 1:**
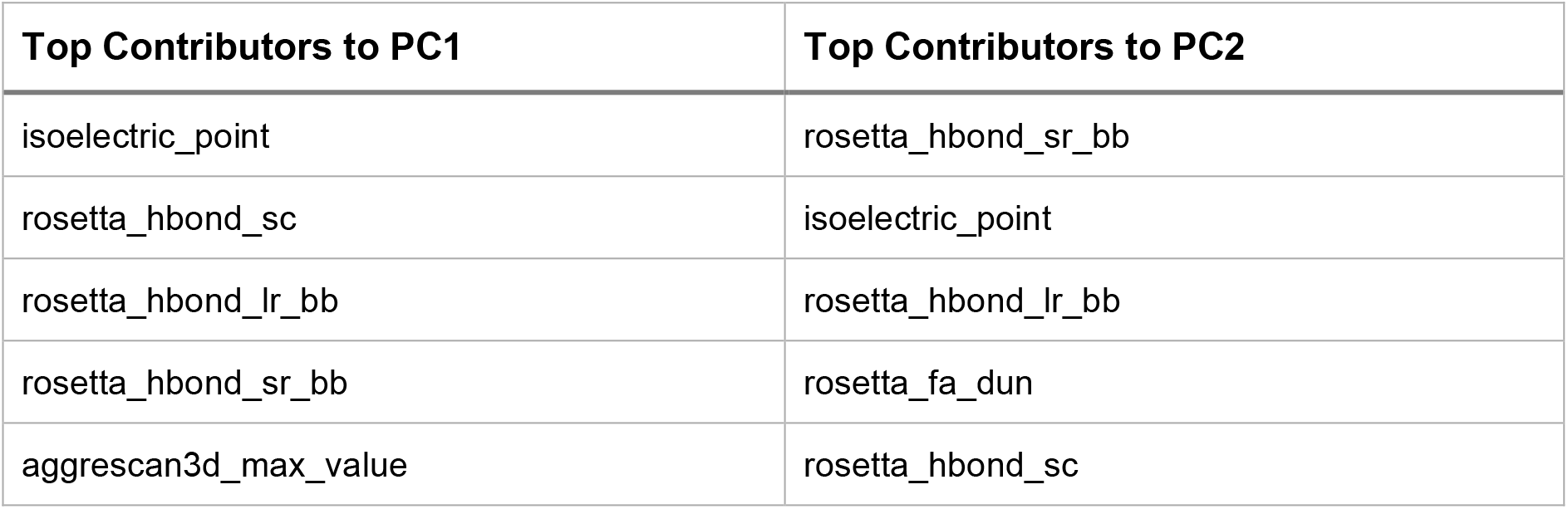
Top 5 contributors to PC1 and PC2 for the AF2 structural model PCA space in figure 3A. For this PCA space the minmax scaling method was used and amino acid composition metrics were excluded.

| Top Contributors to PC1 | Top Contributors to PC2 |
| --- | --- |
| isoelectric_point | rosetta_hbond_sr_bb |
| rosetta_hbond_sc | isoelectric_point |
| rosetta_hbond_lr_bb | rosetta_hbond_lr_bb |
| rosetta_hbond_sr_bb | rosetta_fa_dun |
| aggrescan3d_max_value | rosetta_hbond_sc |

**Figure 3.**
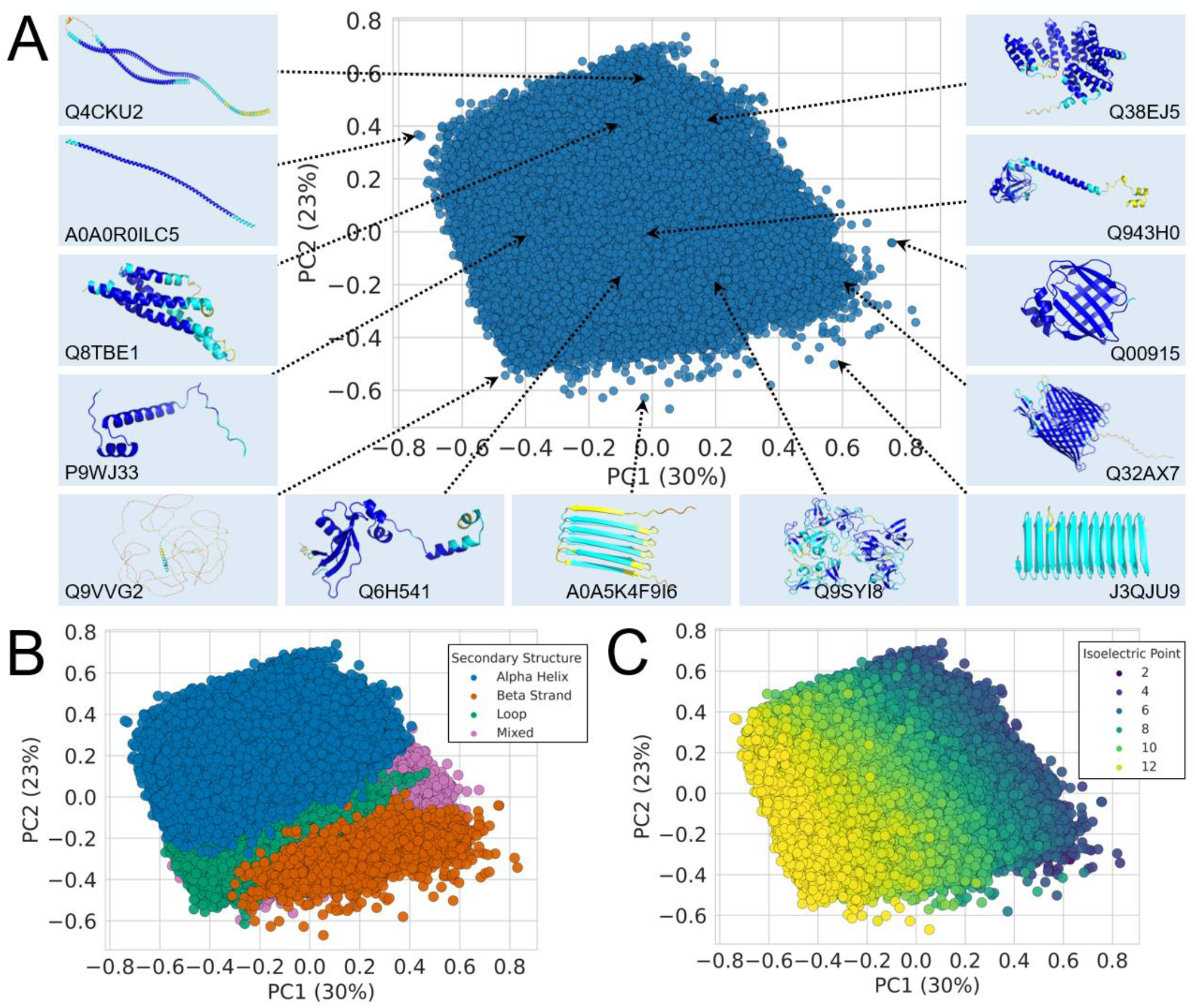
Large-scale analysis of structural features of AF2 models. Amino acid composition metrics are excluded from this analysis. A) A plot of the top two principal components for 564,446 AlphaFold2 predicted structures from 48 model organisms, with a sample of proteins labelled around the PCA space, along with their Uniprot ID and the cartoon representation of the structure. B) The same PCA space as A) but coloured by secondary structure group. C) The same PCA space as A) but coloured by isoelectric point.

### Physicochemical Properties Distinguish Eukaryotic and Prokaryotic Organisms

After observing that simple metrics such as secondary structure composition and isoelectric point are major factors that vary across large structural data sets, we then explored whether there was systematic variation in the physicochemical properties of proteins between organisms. This analysis was performed on a culled data set, which excluded homologous proteins for each organism using MMSeqs2 (Steinegger & Söding, 2017), to remove potential bias from sequence redundancy. In addition to this, we excluded low quality AF2 structural models, with average pLDDT score < 70. Table S21 shows the number of proteins by organism, that were downloaded from AlphaFold DB, the number after filtering for redundancy and the number after removing low quality models.

For each of these data sets, we calculated the DE-STRESS metrics for all proteins, once again excluding amino acid composition, and applied PCA to the average properties per organism (Figure 4A). From this plot it is clear that eukaryotic and prokaryotic proteins are distinct in this space. We also performed similar analysis by applying K-means clustering (MacQueen, 1967) to the average physicochemical properties and found that 2 clusters have the highest mean adjusted rand index (Rand, 1971), supporting this conclusion (Figure 4B). The main contributors to PC1 included Rosetta energy terms capturing rotamer preferences, electrostatics, and Aggrescan3D aggregation propensity scores, while Rosetta energy terms capturing solvation energy and electrostatics, were found to be the main contributors to PC2. The standard scaling method was used for this analysis; however, similar results were found using the minmax and robust scaling methods shown in figure S11.

**Figure 4.**
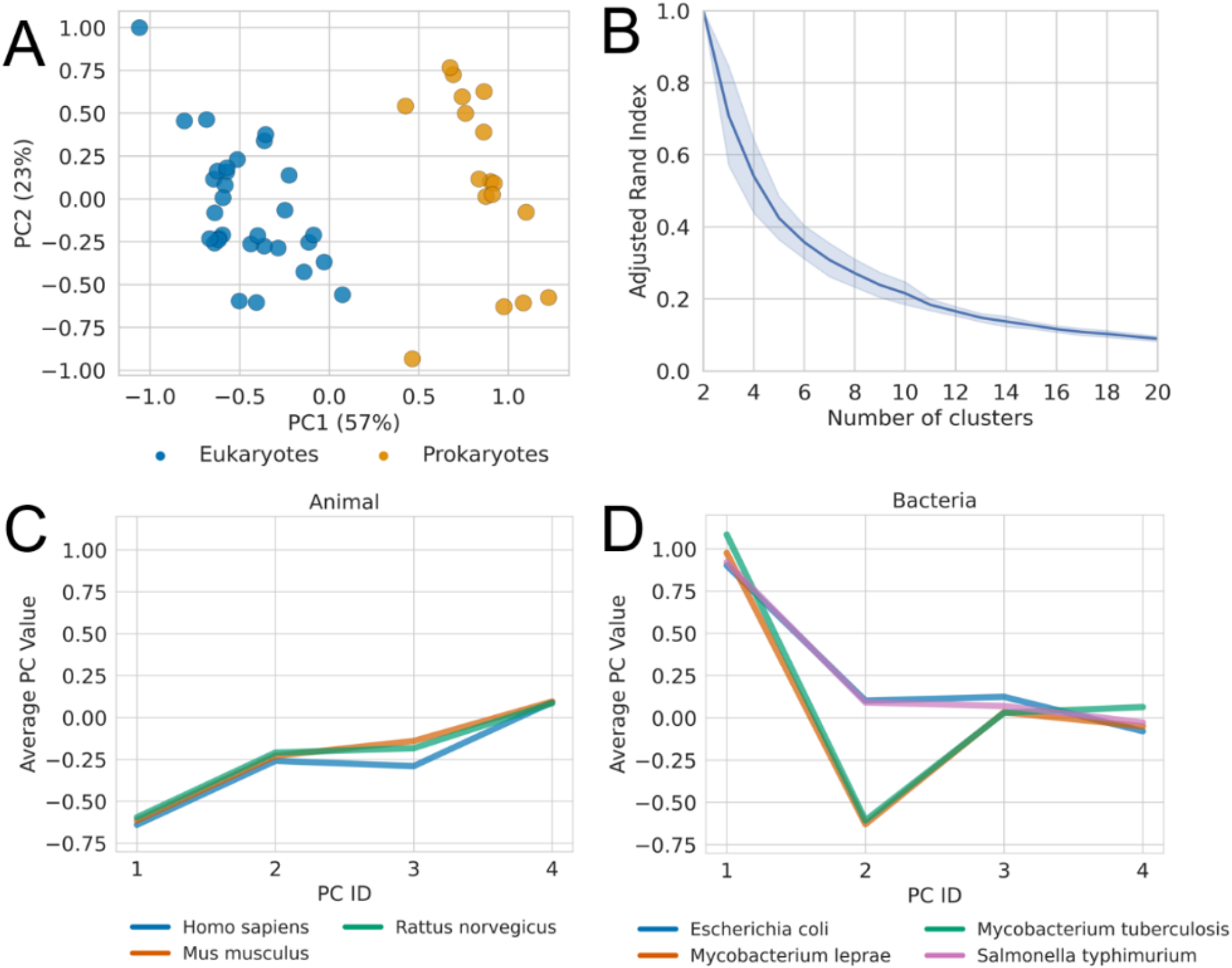
A) PCA analysis of the average model-derived properties for each organism, along with the variance explained. B) The mean and standard deviation of the adjusted rand index against the eukaryote and prokaryote groups, for 100 random initialisations of K-means, and different numbers of clusters. C) The average value of 4 principal components with a 95% confidence interval for 3 different animals. D) The average value of 4 principal components with a 95% confidence interval for 4 different bacteria.

As we determined that eukaryotes and prokaryotes can be distinguished from the average properties of their proteins, we decided to explore this phenomenon at the organism level. Figure 4C&D show the results of applying PCA to the physicochemical properties of all the AF2 structural models from 48 organisms (with homologous proteins and structural models with pLDDT < 70 removed) and plotting the average principal component (PC) values by organism across the animal and bacteria kingdoms. We compared the profiles of 4 PCs, as together they explained over 95% of the variance across the data set. There were clear differences between the profiles, which seemed to be linked to organism type. For example, . *musculus, R. norvegicus* and *H. sapiens* were shown to have extremely similar profiles (Figure 4C) but varied significantly when compared to 4 species of bacteria (Figure 4D). While *E. coli* and *S. typhimurium* shared similar profiles, as did *M. leprae* and *M. tuberculosis*, there was a large difference between these 2 groups of bacteria, indicating that the average properties of their respective proteins are significantly different.

### Reconstructing The Tree of Life from Physicochemical Properties

Following on from the observation that these physicochemical properties can be used to distinguish between individual organisms, a larger scale analysis was performed across all 48 organisms in the data set. Hierarchical agglomerative clustering (Johnson, 1967) was performed on the average physicochemical properties of the organism’s proteomes, using different scaling and linkage methods. We found that the clustering results largely recreated the tree of life and captured relationships observed at a DNA sequence level from structural properties of protein models alone (Figure 5). After this, we compared these trees to a reference tree from NCBI (Schoch et al., 2020), using the clustering information distance (Smith, 2020). For context, a distance of 0 is an exact match and the expected value for 1000 randomly generated trees with 48 leaf nodes was found to be 0.89 (Smith, 2020). The highest ranked tree had a clustering information distance of 0.43; however, the other trees had comparable distances (Table S20). This indicates significant similarities between the trees.

**Figure 5.**
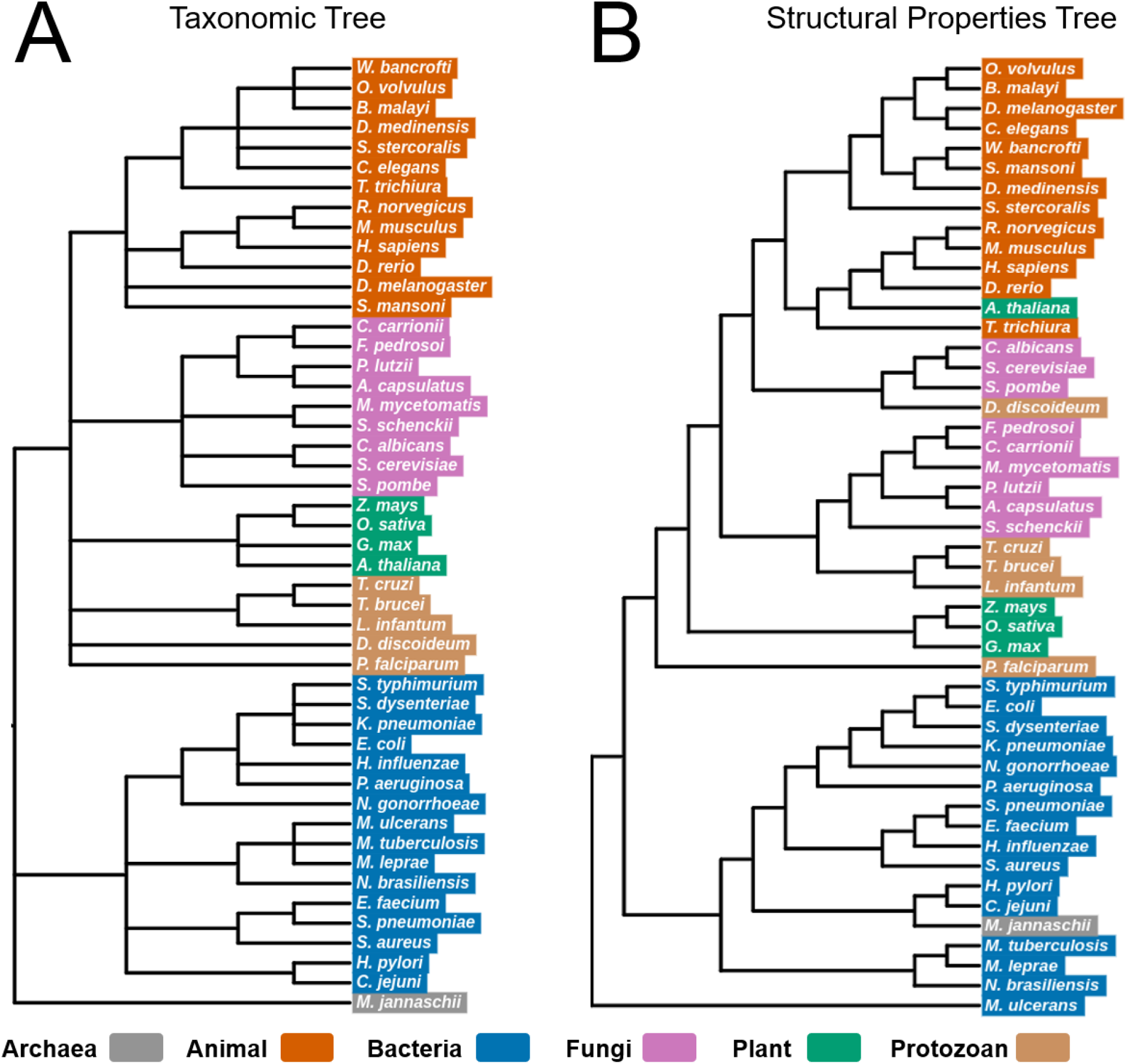
A) A tree of the 48 organisms created from the Common Tree tool from the NCBI Taxonomy Browser, which is based on a diverse set of phylogenetic information. B) The highest ranked tree created from hierarchical clustering on the average model-derived properties of each organism.

Qualitatively, there are broad similarities between the trees, such as the separation of prokaryotes and eukaryotes, consistent with figure 4A&B, as well as details that make sense, such as clusters composed of similar organisms: the mammals *R. norvegicus, M. musculus* and *H. sapiens*; the fungi *P. lutzii* and *A. capsulatus*; the crop plants *Z. mays, O. sativa* and *G. max*; and the pathogenic enteric bacteria *E. coli* and *S. typhimurium*. However, there are also interesting differences. For example, the NCBI taxonomic tree has three main clusters, consisting of eukaryotes, bacteria and archaea, while the archaea *M. jannaschii* is grouped together with bacteria in the structural properties tree, despite this organism being a thermophilic methanogen (Jones et al., 1983) that experiences a very different cellular environment. Furthermore, in the structural properties tree, the bacteria *M. ulcerans* is in a cluster on its own, separated from the rest of the bacteria. *M. ulcerans* is a pathogenic bacterium that has evolved to live in a restricted environment (Doig et al., 2012) and has undergone significant genome reduction (Demangel et al., 2009), which is likely to have impacted the average properties of its proteins. Another difference observed in the structural properties tree is that the protozoan *D. discoideum*, groups together with the fungi, rather than the other protozoans. *D. discoideum* has a complex life cycle, that begins as a single-celled amoeba, which transforms into a multi-cellular slug, and finally it becomes a fruiting body that releases spores (Fey et al., 2007). This complex life cycle might differentiate it from other protozoans. Curiously, one other difference observed from this tree is that *A. thaliana* groups together with animals such as *D. rerio* and *T. trichiura*. It is not entirely obvious why this could be the case; however, *A. thaliana* does have a much smaller proteome and a shorter life cycle than the other plants included in this analysis (Carneiro et al., 2015; Müller & Grossniklaus, 2010).

## Discussion

With the development of novel structure prediction algorithms, we now have access to structural data that was previously unavailable, enabling us to apply a range of data analysis techniques to gain new insights into protein structure and function. We have demonstrated that physicochemical properties derived from predicted structures are useful on small datasets and can be predictive of protein production of designed antibodies using only simple methods that requires little to no fitting of hyperparameters. Key structural features identified in this data-driven approach (Tables S12 and S13), aligned with factors that the designers explicitly discussed as being linked with low levels of expression during experimental characterisation of the designs, such as cavities in the core of the structure (hydrophobic_fitness), unpaired buried charges (rosetta_fa_elec) and loss of long range hydrogen bonds (rosetta_hbond_bb_sc) (Baran et al., 2017). In addition to this, aggregation propensity of designs (aggrescan3d_avg_value, aggrescan3d_min_value, aggrescan3d_max_value), were also shown to be important for separating out low and high producing designs, which makes sense as aggregation commonly causes low production of designed proteins (P. S. Huang et al., 2016). Furthermore, these results appear to generalise beyond the initial set of designed scFv antibodies to other scFvs that have been experimentally characterised from the Structural Antibody Database (SAbDab), and so this could form the basis of a method to optimise protein production for applications in biotechnology.

At scale, these structural physicochemical descriptors can be used to uncover different kinds of relationships. Our analysis of over half a million AlphaFold2 (AF2) structural models from 48 different organisms, demonstrated that one of the biggest differences between proteins related to secondary structure. This satisfyingly supports decades of efforts to classify proteins by fold, such as in the CATH (Bordin et al., 2023) and SCOP (Andreeva et al., 2014, 2020) databases. We also saw similar relationships when analysing 160,000 experimentally determined structures from the Protein Data Bank (PDB) (Figure S7), which suggest that these properties vary across protein structures in general, rather than just AF2 structural models.

Next, we analysed the average properties of proteins by organism, to determine if there was systematic variation. We found that eukaryotic and prokaryotic organisms were readily separable, which is likely to be the result of known differences between eukaryotic and prokaryotic proteins, such as differences in the level of disordered proteins (Basile et al., 2019), multi-domain proteins (Brocchieri & Karlin, 2005), and isoelectric point (Kiraga et al., 2007). Eukaryotic and prokaryotic organisms are mainly separated across principal component 1, which has Rosetta energy terms capturing rotamer preferences, electrostatics and Aggrescan3D aggregation propensity scores as the main contributors. Previous research has found statistically significant differences in rotamer preferences between trans-membrane and soluble proteins (Chamberlain & Bowie, 2004), which may explain this variance observed between eukaryotes and prokaryotes, as there could be differences in the proportions of trans-membrane and soluble proteins across these organisms.

Lastly, we found that even individual organisms can be distinguished by the average physicochemical properties of their proteins. Initially, this was shown in figure 4C&D for a subset of animal and bacteria organisms, as we see that *S. typhimurium* and *E. coli* are both separated from *M. leprae* and *M. tuberculosis* across PC2. Rosetta energy terms capturing solvation energy and electrostatics, are the major factors that contribute to principal component 2, which could be an indicator of the differences in environments in which these organisms have evolved. After observing these relationships, we then explored this across all 48 organisms in the data set using hierarchical agglomerative clustering. Remarkably, the trees created from the average physicochemical properties of proteins, are largely consistent with a tree generated from diverse phylogenetic information from NCBI, such as DNA sequence information. This is surprising, as the chemistry of proteins is functionally constrained to a much greater extent than DNA sequences (Zhang & Yang, 2015).

As this type of analysis has only become possible over the last few years, this is, to our knowledge, the first time these relationships have been demonstrated. Since amino acid and secondary structure composition metrics were removed from the data set for the large-scale clustering analysis, the clustering is performed only on structure-derived properties of the proteins. These results indicate that the properties of proteins could be optimised to the chemical environment that the protein will experience, something that has been largely ignored to date, while engineering and designing proteins. This observation could lead to the development of more robust design methodologies that incorporate this information and increase the reliability of protein design and engineering in the future.

## Methods

### Datasets

#### Fleishman and SAbDab Scfv Antibody Fragment Data Sets

The Fleishman data set consists of 192 single chain variable fragment (scFv) amino acid sequences, along with a yeast display protein production measure (Baran et al., 2017). Five unrelaxed structural models of the scFv sequences were obtained using AlphaFold2 (AF2) (Jumper et al., 2021), and the highest ranked model was selected for each design. In addition to this, a set of non-redundant, experimentally determined scFvs were obtained from the Structural Antibody Database (SAbDab) (Dunbar et al., 2014; Schneider et al., 2022). This database contains all the antibody structures from the Protein Data Bank (PDB) (Berman et al., 2000), and it was used to search for scFvs that had a maximum sequence identity of 90%, were not in a complex with an antigen, and included both heavy and light chain regions. After obtaining this data set of 41 experimentally determined scFvs, AF2 was then used to generate unrelaxed, structural models for these scFvs, in order to be consistent with the Fleishman data set. This is because there could be inherent differences between the SAbDab structures and the AF2 structural models from biases in the AF2 structure prediction. This is important as this could confound our analysis of the physicochemical properties and scFv protein production.

#### AlphaFold DB and PDB Data Sets

AF2 structural models for model organisms and global health proteomes, were obtained from the AlphaFold Protein Structural Database (Varadi et al., 2022) https://alphafold.ebi.ac.uk/download. In total, there were 564,446 structural models for 48 organisms, including animals, archaea, bacteria, fungi, plants and protozoans, which were downloaded as PDB files. Table S21 in the supplementary materials shows the full list of 48 organisms, along with the corresponding volume of protein structural models in the data set. For the PDB data set, a total of 189,942 experimentally determined protein structures were downloaded from the Protein Data Bank as PDB files. This data set is up to date as of July 2020.

### Data preparation

#### Fleishman and SAbDab Scfv Antibody Fragment Data Sets

Our model evaluation software DE-STRESS (Stam & Wood, 2021) was used to generate physicochemical properties from the structural models in the Fleishman and SAbDab scFv data sets, and these were used as the features throughout this analysis. These properties included all-atom scoring functions (Alford et al., 2017; X. Huang et al., 2020; McIntosh-Smith et al., 2012, 2015; Yang & Zhou, 2008), geometric metrics such as packing density (MS, 2007; Wood et al., 2017) and hydrophobic fitness (E. S. Huang et al., 1995; Wood et al., 2017), aggregation propensity (Kuriata et al., 2019), isoelectric point, and amino acid and secondary structure composition. All these metrics were generated from structural models of folded proteins and did not include any information on DNA or mRNA sequences.

Author reported protein production values, which were found from performing yeast display experiments on the scFv designs (Baran et al., 2017), were split into three equal classes: low, medium and high. These three classes were informed by analysing the distribution of protein production values across the whole data set. Histograms of these values showed peaks at the low and high protein production levels, and lower volumes around the medium production levels. After this, these protein production classes were used for stratified sampling, in order to create a 75% training set and 25% test set.

Several different data processing, feature selection and scaling methods were applied to the training set to prepare it for analysis and model training. All-atom scoring functions and hydrophobic fitness values were normalised by the number of residues in each design, as sequence length impacts the magnitude of these energy value features. In addition to this, features that were constant or had the same value for more than 75% of the samples in the training set, were identified and removed, and the data set was scaled using the standard, robust and minmax scaling methods. Highly correlated features were determined using the spearman correlation coefficient (Spearman, 1987) (0.7 or higher), and these features were removed sequentially until no correlated features remained in the data set. All these steps were repeated, including and excluding the amino acid composition metrics, and with two different methods of feature selection. One of these methods involved calculating the mutual information score (Mackay, 2003) of each feature against the protein production classes, and retaining those features that had a score greater than the mean mutual information score of all features. In addition to this, a random forest (Breiman, 2001) was used with the number of estimators set to 1000 and a balanced class weight. Finally, the scalers used, and features removed from the training set, were also applied to the SAbDab and test scFv data sets as well.

#### AlphaFold DB and PDB Data Sets

Firstly, 564,446 PDB files were downloaded from the AlphaFold DB and the DE-STRESS metrics were calculated for these files, using the headless version of the DE-STRESS software. Features that contained greater than 5% missing values were dropped from the dataset, along with amino acid and secondary structure composition metrics. Once these features had been dropped, any rows that had missing values for the rest of the features were removed, which resulted in 564,432 structural models left in the data set. Next, the Uniprot API (Consortium, 2023) was used to extract additional information about each of the protein structures in the data set, including the organism that the protein originates from.

This information was then joined onto the data set of DE-STRESS metrics by Uniprot id, and any duplicates in the data set were removed. After this, constant features were dropped, the data set was scaled with the minmax, standard and robust methods, and highly correlated features were removed in the same way as the Fleishman scFv data set. This data set was used for the analysis in figure 3; however, for the organism clustering in figure 4 and 5, we also excluded homologous sequences in order to remove bias. MMseqs2 (Steinegger & Söding, 2017) easy cluster with min_seq_id set to 0.3, was used to remove these sequences. After this, the output was used to filter the data set to 387,810 non-redundant proteins. In addition to this, low quality AF2 structural models were excluded, by removing models with average pLDDT < 70%, which resulted in 241,134 structural models in the organism clustering. The final set of DE-STRESS metrics used in this data set were isoelectric_point, budeff_charge, evoef2_ref_total, rosetta_lk_ball_wtd, rosetta_fa_intra_sol_xover4, rosetta_hbond_lr_bb, rosetta_hbond_sr_bb, rosetta_hbond_sc, rosetta_fa_dun, aggrescan3d_avg_value, aggrescan3d_min_value, aggrescan3d_max_value.

Next, 189,942 structures from the PDB were processed in the same way as the AF2 structural models, except the MMseqs2 step was not used in this case. As the PDB data set is smaller than the AF2 data set, a threshold of 20% was used to remove features that had a lot of missing values. After removing rows that still had missing values remaining across the features, there were 165,293 structures left in the data set. Designed proteins were identified and labelled in this data set using a curated list (Woolfson, 2021), and the rest of the PDB structures were labelled as “native”. The final set of DE-STRESS metrics used in this data set were; isoelectric_point, packing_density, evoef2_ref_total, rosetta_fa_elec, rosetta_fa_sol, rosetta_lk_ball_wtd, rosetta_fa_intra_sol_xover4, rosetta_hbond_bb_sc, rosetta_rama_prepro, rosetta_p_aa_pp, rosetta_fa_dun, aggrescan3d_avg_value, aggrescan3d_min_value, aggrescan3d_max_value.

### Predicting Protein Production from Physicochemical Properties Of Proteins

Principal component analysis (PCA) (Pearson, 1901) was used to explore whether the physicochemical properties could distinguish scFv designs with low and high protein production. This was performed on the different training sets with the minmax, standard and robust scaling methods, including and excluding amino acid composition metrics, and using a random forest (Breiman, 2001) and mutual information (Mackay, 2003) for feature selection. The features that contributed to the principal components (PCs) were calculated along with the variance explained by each component. The AF2 structural models of the 41 SAbDab scFvs were also included in this analysis, to see how they compared to the designed scFvs across the physicochemical properties. After performing PCA, stratified 5-fold cross validation with 10 random splits was used to train and validate Naive Bayes classifiers (Bishop, 2006) on the different training sets. These models were evaluated with weighted precision, recall, one-vs-rest multiclass ROC curves, and confusion matrices. Finally, these models were also evaluated on the 25% test set with the same metrics.

### Large-Scale Analysis of Physicochemical Properties Performed Across Half A Million Predicted Protein Structures

PCA was applied to the data sets of 564,432 AF2 structural models and 165,293 PDB structures, scaled using the minmax and robust scaling methods. Scatter plots of PC1 and PC2 were created, and a sample of proteins was labelled around these spaces. In addition to this, the main contributors to PC1 and PC2 were calculated and plots were created to show how metrics such as secondary structure composition, isoelectric point, packing density and aggregation propensity, varied across this space. This was shown across the scatter plots of PC1 and PC2, but also as cumulative histograms.

### Exploring The Relationships Between Organisms with Physicochemical Properties

For all three scaling methods, the average physicochemical properties were computed for the 48 organisms, and then PCA and clustering methods were applied to these data sets. K-means (MacQueen, 1967) was performed with 100 random initialisations, with the number of clusters ranging between 2 and 20 clusters. The adjusted rand index (Rand, 1971) was used to compare the clustering labels against labels indicating whether the organism was eukaryotic or prokaryotic. Hierarchical agglomerative clustering (Johnson, 1967) was then performed on the average physicochemical properties of each organism, with the Euclidean distance metric and four different linkage methods: single, average, complete and ward.

Dendrograms were created for each of these clusterings and then compared to a tree of the 48 organisms created from the common tree tool from the NCBI Taxonomy Browser (Schoch et al., 2020). The clustering information distance metric (Smith, 2020) was used to compare each of the dendrograms of the 48 organisms against the NCBI tree. Finally, the interactive tree of life online tool (Letunic & Bork, 2021) was used to display these trees and to sort the trees by the number of leaf nodes, with the fewest at the bottom and the most at the top.

## Funding

CWW is supported by an Engineering and Physical Sciences Research Council Fellowship (EP/S003002/1). Michael Stam is supported by the United Kingdom Research and Innovation (grant EP/S02431X/1), UKRI Centre for Doctoral Training in Biomedical AI at the University of Edinburgh, School of Informatics. This work was supported by the Wellcome Trust-University of Edinburgh Institutional Strategic Support Fund (ISSF3).

## Notes

### Competing Interest Statement

The authors have declared no competing interest.

