## Supplementary Material for "Large scale analysis of predicted protein structures links model features to *in vivo* behaviour"

### Supplementary Materials

#### Glossary of DE-STRESS metrics

| Metric Name | Metric Description | Variable Name in CSV Output |
|---|---|---|
| Total Score | This value is a global indicator of the aggregation propensity/solubility of the protein structure. It depends on the protein size. It allows assessing changes in solubility promoted by amino acid substitutions in a particular protein structure. The more negative the value, the highest the global solubility. | aggrescan3d_total_value |
| Average Score | This value is a normalized indicator of the aggregation propensity/solubility of the protein structure. Allows comparing the solubility of different protein structures. It also allows assessing changes in solubility promoted by amino acid substitutions in a particular protein structure. The more negative the value, the highest the normalized solubility. | aggrescan3d_avg_value |
| Minimum Score | This is the value of the most soluble residue in the structural context. | aggrescan3d_min_value |
| Maximum Score | This is the value of the most aggregation-prone residue in the structural context. | aggrescan3d_max_value |

*Table S1: This table provides a description of the metrics from the Aggrescan3D 2.0 programme and their variable names in the CSV output file.*

| Metric Name | Metric Description | Variable Name in CSV Output |
|---|---|---|
| Total Energy | This value is the total BUDE force field energy. It is the sum of the steric, desolvation and charge components. | budeff_total |
| Steric Energy | This value is the steric component of the BUDE force field energy. It is calculated with a simplified Leonard-Jones potential. It is softer than the steric component of many other force fields. | budeff_steric |
| Desolvation Energy | This value is the desolvation component of the BUDE force field energy. | budeff_desolvation |
| Charge Energy | This value is the charge component of the BUDE force field energy. | budeff_charge |

*Table S2: This table provides a description of the metrics from the BUDE programme and their variable names in the CSV output file.*

| Metric Name | Metric Description | Variable Name in CSV Output |
|---|---|---|
| Total Energy | This value is the total DFIRE2 energy. This is the only field that is returned from running DFIRE2 on a pdb file. | dfire2_total |

*Table S3: This table provides a description of the metrics from the DFIRE2 programme and their variable names in the CSV output file.*

| Metric Name | Metric Description | Variable Name in CSV Output |
|---|---|---|
| H | α-helix | ss_prop_alpha_helix |
| B | Isolated β-bridge | ss_prop_beta_bridge |
| E | Extended β-strand | ss_prop_beta_strand |
| G | 3-10 helix | ss_prop_3_10_helix |
| I | π-helix | ss_prop_pi_helix |
| T | Hydrogen-bonded turn | ss_prop_hbonded_turn |
| S | Bend | ss_prop_bend |
| - | Loop | ss_prop_loop |

*Table S4: This table provides a description of the metrics from the DSSP programme and their variable names in the CSV output file.*

| Metric Name | Metric Description | Variable Name in CSV Output |
|---|---|---|
| Total Energy | This value is the total EvoEF2 energy. It is the sum of the reference, intra residue, inter residue - same chain and inter residue - different chains, energy values. In the EvoEF2 output this field is called `Total`. | evoef2_total |
| Reference Energy | This value is the total reference energy. This value is not included in the EvoEF2 output and is calculated in DE-STRESS. | evoef2_ref_total |
| Intra Residue Energy | This value is the total energy for intra residue interactions. This value is not included in the EvoEF2 output and is calculated in DE-STRESS. | evoef2_intraR_total |
| Inter Residue - Same Chain Energy | This value is the total energy for inter residue interactions in the same chain. This value is not included in the EvoEF2 output and is calculated in DE-STRESS. | evoef2_interS_total |
| Inter Residue - Different Chains Energy | This value is the total energy for inter residue interactions in different chains. This value is not included in the EvoEF2 output and is calculated in DE-STRESS. | evoef2_interD_total |
| ALA - Reference | This value is reference energy for the amino acid Alanine (ALA). In the EvoEF2 output this value is called `reference_ALA`. | Not included in csv output |
| ARG - Reference | This value is reference energy for the amino acid Arginine (ARG). In the EvoEF2 output this value is called `reference_ARG`. | Not included in csv output |
| ASN - Reference | This value is reference energy for the amino acid Asparagine (ASN). In the EvoEF2 output this value is called `reference_ASN`. | Not included in csv output |
| ASP - Reference | This value is reference energy for the amino acid Aspartic acid (ASP). In the EvoEF2 output this value is called `reference_ASP`. | Not included in csv output |
| CYS - Reference | This value is reference energy for the amino acid Cysteine (CYS). In the EvoEF2 output this value is called `reference_CYS`. | Not included in csv output |
| GLN - Reference | This value is reference energy for the amino acid Glutamine (GLN). In the EvoEF2 output this value is called `reference_GLN`. | Not included in csv output |
| GLU - Reference | This value is reference energy for the amino acid Glutamic acid (GLU). In the EvoEF2 output this value is called `reference_GLU`. | Not included in csv output |

| | | |
|---|---|---|
| GLY - Reference | This value is reference energy for the amino acid glycine (GLY). In the EvoEF2 output this value is called `reference_GLY`. | Not included in csv output |
| HIS - Reference | This value is reference energy for the amino acid Histidine (HIS). In the EvoEF2 output this value is called `reference_HIS`. | Not included in csv output |
| ILE - Reference | This value is reference energy for the amino acid Isoleucine (ILE). In the EvoEF2 output this value is called `reference_ILE`. | Not included in csv output |
| LEU - Reference | This value is reference energy for the amino acid Leucine (LEU). In the EvoEF2 output this value is called `reference_LEU`. | Not included in csv output |
| LYS - Reference | This value is reference energy for the amino acid Lysine (LYS). In the EvoEF2 output this value is called `reference_LYS`. | Not included in csv output |
| MET - Reference | This value is reference energy for the amino acid Methionine (MET). In the EvoEF2 output this value is called `reference_MET`. | Not included in csv output |
| PHE - Reference | This value is reference energy for the amino acid Phenylalanine (PHE). In the EvoEF2 output this value is called `reference_PHE`. | Not included in csv output |
| PRO - Reference | This value is reference energy for the amino acid Proline (PRO). In the EvoEF2 output this value is called `reference_PRO`. | Not included in csv output |
| SER - Reference | This value is reference energy for the amino acid Serine (SER). In the EvoEF2 output this value is called `reference_SER`. | Not included in csv output |
| THR - Reference | This value is reference energy for the amino acid Threonine (THR). In the EvoEF2 output this value is called `reference_THR`. | Not included in csv output |
| TRP - Reference | This value is reference energy for the amino acid Tryptophan (TRP). In the EvoEF2 output this value is called `reference_TRP`. | Not included in csv output |
| TYR - Reference | This value is reference energy for the amino acid Tyrosine (TYR). In the EvoEF2 output this value is called `reference_TYR`. | Not included in csv output |
| VAL - Reference | This value is reference energy for the amino acid Valine (VAL). In the EvoEF2 output this value is called `reference_VAL`. | Not included in csv output |
| VDW Attractive - Intra Residue | This value is the Van der Waals attractive energy for intra residue interactions. In the EvoEF2 output this value is called `intraR_vdwatt`. | Not included in csv output |
| VDW Repulsive - Intra Residue | This value is the Van der Waals repulsive energy for intra residue interactions. In the EvoEF2 output this value is called `intraR_vdwrep`. | Not included in csv output |
| Electrostatics - Intra Residue | This value is the Coulomb's electrostatics energy for intra residue interactions. In the EvoEF2 output this value is called `intraR_electr`. | Not included in csv output |
| Desolvation Polar - Intra Residue | This value is the polar atoms desolvation energy for intra residue interactions. In the EvoEF2 output this value is called `intraR_deslvP`. | Not included in csv output |
| Desolvation Non Polar - Intra Residue | This value is the non polar atoms desolvation energy for intra residue interactions. In the EvoEF2 output this value is called `intraR_deslvH`. | Not included in csv output |
| HB Sidechain Backbone Distance - Intra Residue | This value is the energy for the hydrogen-acceptor distance from sidechain - backbone and intra residue interactions. In the EvoEF2 output this value is called `intraR_hbscbb_dis`. | Not included in csv output |

| | | |
|---|---|---|
| HB Sidechain Backbone Theta - Intra Residue | This value is the energy for the angle between the donor, hydrogen and acceptor atoms (theta), from sidechain - backbone and intra residue interactions. In the EvoEF2 output this value is called `intraR_hbscbb_the`. | Not included in csv output |
| HB Sidechain Backbone Phi - Intra Residue | This value is the energy for the angle between the hydrogen, acceptor and base atoms (phi), from sidechain - backbone and intra residue interactions. In the EvoEF2 output this value is called `intraR_hbscbb_phi`. | Not included in csv output |
| Amino Acid Propensity - Intra Residue | This value is the amino acid propensity energy for intra residue interactions. In the EvoEF2 output this value is called `aapropensity`. | Not included in csv output |
| Ramachandran - Intra Residue | This value is the Ramachandran energy for intra residue interactions. In the EvoEF2 output this value is called `ramachandran`. | Not included in csv output |
| Dunbrack Rotamer - Intra Residue | This value is the Dunbrack Rotamer energy for intra residue interactions. In the EvoEF2 output this value is called `dunbrack`. | Not included in csv output |
| VDW Attractive - Inter Residue - Same Chain | This value is the Van der Waals attractive energy for inter residue interactions - same chain. In the EvoEF2 output this value is called `interS_vdwatt`. | Not included in csv output |
| VDW Repulsive - Inter Residue - Same Chain | This value is the Van der Waals repulsive energy for inter residue interactions - same chain. In the EvoEF2 output this value is called `interS_vdwrep`. | Not included in csv output |
| Electrostatics - Inter Residue - Same Chain | This value is the Coulomb's electrostatics energy for inter residue interactions - same chain. In the EvoEF2 output this value is called `interS_electr`. | Not included in csv output |
| Desolvation Polar - Inter Residue - Same Chain | This value is the polar atoms desolvation energy for inter residue interactions - same chain. In the EvoEF2 output this value is called `interS_deslvP`. | Not included in csv output |
| Desolvation Non Polar - Inter Residue - Same Chain | This value is the non polar atoms desolvation energy for inter residue interactions - same chain. In the EvoEF2 output this value is called `interS_deslvH`. | Not included in csv output |
| Disulfide Bonding - Inter Residue - Same Chain | This value is the disulfide bonding energy for inter residue interactions - same chain. In the EvoEF2 output this value is called `interS_ssbond`. | Not included in csv output |
| HB Backbone Backbone Distance - Inter Residue - Same Chain | This value is the energy for the hydrogen-acceptor distance from backbone - backbone and inter residue interactions - same chain. In the EvoEF2 output this value is called `interS_hbbbbb_dis`. | Not included in csv output |
| HB Backbone Backbone Theta - Inter Residue - Same Chain | This value is the energy for the angle between the donor, hydrogen and acceptor atoms (theta), from backbone - backbone and inter residue interactions - same chain. In the EvoEF2 output this value is called `interS_hbbbbb_the`. | Not included in csv output |
| HB Backbone Backbone Phi - Inter Residue - Same Chain | This value is the energy for the angle between the hydrogen, acceptor and base atoms (phi), from backbone - backbone and inter residue interactions - same chain. In the EvoEF2 output this value is called `interS_hbbbbb_phi`. | Not included in csv output |
| HB Sidechain Backbone Distance - Inter Residue - Same Chain | This value is the energy for the hydrogen-acceptor distance from side chain - backbone and inter residue interactions - same chain. In the EvoEF2 output this value is called `interS_hbscbb_dis`. | Not included in csv output |
| HB Sidechain Backbone Theta - Inter Residue - Same Chain | This value is the energy for the angle between the donor, hydrogen and acceptor atoms (theta), from side chain - backbone and inter residue interactions - same chain. In the EvoEF2 output this value is called `interS_hbscbb_the`. | Not included in csv output |

| | | |
|---|---|---|
| HB Sidechain Backbone Phi - Inter Residue - Same Chain | This value is the energy for the angle between the hydrogen, acceptor and base atoms (phi), from side chain - backbone and inter residue interactions - same chain. In the EvoEF2 output this value is called `interS_hbscbb_phi`. | Not included in csv output |
| HB Sidechain Sidechain Distance - Inter Residue - Same Chain | This value is the energy for the hydrogen-acceptor distance from side chain - side chain and inter residue interactions - same chain. In the EvoEF2 output this value is called `interS_hbscsc_dis`. | Not included in csv output |
| HB Sidechain Sidechain Theta - Inter Residue - Same Chain | This value is the energy for the angle between the donor, hydrogen and acceptor atoms (theta), from side chain - side chain and inter residue interactions - same chain. In the EvoEF2 output this value is called `interS_hbscsc_the`. | Not included in csv output |
| HB Sidechain Sidechain Phi - Inter Residue - Same Chain | This value is the energy for the angle between the hydrogen, acceptor and base atoms (phi), from side chain - side chain and inter residue interactions - same chain. In the EvoEF2 output this value is called `interS_hbscsc_phi`. | Not included in csv output |
| VDW Attractive - Inter Residue - Different Chains | This value is the Van der Waals attractive energy for inter residue interactions - different chains. In the EvoEF2 output this value is called `interD_vdwatt`. | Not included in csv output |
| VDW Repulsive - Inter Residue - Different Chains | This value is the Van der Waals repulsive energy for inter residue interactions - different chains. In the EvoEF2 output this value is called `interD_vdwrep`. | Not included in csv output |
| Electrostatics - Inter Residue - Different Chains | This value is the Coulomb's electrostatics energy for inter residue interactions - different chains. In the EvoEF2 output this value is called `interD_electr`. | Not included in csv output |
| Desolvation Polar - Inter Residue - Different Chains | This value is the polar atoms desolvation energy for inter residue interactions - different chains. In the EvoEF2 output this value is called `interD_deslvP`. | Not included in csv output |
| Desolvation Non Polar - Inter Residue - Different Chains | This value is the non polar atoms desolvation energy for inter residue interactions - different chains. In the EvoEF2 output this value is called `interD_deslvH`. | Not included in csv output |
| Disulfide Bonding - Inter Residue - Different Chains | This value is the disulfide bonding energy for inter residue interactions - different chains. In the EvoEF2 output this value is called `interD_ssbond`. | Not included in csv output |
| HB Backbone Backbone Distance - Inter Residue - Different Chains | This value is the energy for the hydrogen-acceptor distance from backbone - backbone and inter residue interactions - different chains. In the EvoEF2 output this value is called `interD_hbbbbb_dis`. | Not included in csv output |
| HB Backbone Backbone Theta - Inter Residue - Different Chains | This value is the energy for the angle between the donor, hydrogen and acceptor atoms (theta), from backbone - backbone and inter residue interactions - different chains. In the EvoEF2 output this value is called `interD_hbbbbb_the`. | Not included in csv output |
| HB Backbone Backbone Phi - Inter Residue - Different Chains | This value is the energy for the angle between the hydrogen, acceptor and base atoms (phi), from backbone - backbone and inter residue interactions - different chains. In the EvoEF2 output this value is called `interD_hbbbbb_phi`. | Not included in csv output |
| HB Sidechain Backbone Distance - Inter Residue - Different Chains | This value is the energy for the hydrogen-acceptor distance from side chain - backbone and inter residue interactions - different chains. In the EvoEF2 output this value is called `interD_hbscbb_dis`. | Not included in csv output |
| HB Sidechain Backbone Theta - Inter Residue - Different Chains | This value is the energy for the angle between the donor, hydrogen and acceptor atoms (theta), from side chain - backbone and inter residue interactions - different chains. | Not included in csv output |

| | | |
|---|---|---|
| | In the EvoEF2 output this value is called `interD_hbscbb_the`. | |
| HB Sidechain Backbone Phi - Inter Residue - Different Chains | This value is the energy for the angle between the hydrogen, acceptor and base atoms (phi), from side chain - backbone and inter residue interactions - different chains. In the EvoEF2 output this value is called `interD_hbscbb_phi`. | Not included in csv output |
| HB Sidechain Sidechain Distance - Inter Residue - Different Chains | This value is the energy for the hydrogen-acceptor distance from side chain - side chain and inter residue interactions - different chains. In the EvoEF2 output this value is called `interD_hbscsc_dis`. | Not included in csv output |
| HB Sidechain Sidechain Theta - Inter Residue - Different Chains | This value is the energy for the angle between the donor, hydrogen and acceptor atoms (theta), from side chain - side chain and inter residue interactions - different chains. In the EvoEF2 output this value is called `interD_hbscsc_the`. | Not included in csv output |
| HB Sidechain Sidechain Phi - Inter Residue - Different Chains | This value is the energy for the angle between the hydrogen, acceptor and base atoms (phi), from side chain - side chain and inter residue interactions - different chains. In the EvoEF2 output this value is called `interD_hbscsc_phi`. | Not included in csv output |

*Table S5: This table provides a description of the metrics from the EvoEF2 programme and their variable names in the CSV output file.*

| **Metric Name** | **Metric Description** | **Variable Name in CSV Output** |
|---|---|---|
| Charge | This value is the total charge of the protein sequence. | charge |
| Hydrophobic Fitness | This value is an efficient centroid-based method for calculating the packing quality of the protein structure. For this method C, F, I, L, M, V, W and Y are considered hydrophobic. | hydrophobic_fitness |
| Isoelectric Point | This value is the pH of a solution at which the net charge of the protein becomes zero. | isoelectric_point |
| Packing Density | This value is the mean packing density of the protein structure, where the packing density of a non-hydrogen atom, is defined as the number of non-hydrogen atoms within a specified radius, for example a radius of 7Å. | packing_density |

*Table S6: This table provides a description of the metrics from the Isambard programme and their variable names in the CSV output file.*

| **Metric Name** | **Metric Description** | **Variable Name in CSV Output** |
|---|---|---|
| Total Energy | This value is the total Rosetta energy. It is a weighted sum of the different Rosetta energy values. In the Rosetta `score.sc` output file, this value is called `total_score`. | rosetta_total |
| Reference | This value is the reference energy for the different amino acids. In the Rosetta `score.sc` output file, this value is called `ref`. | Not included in csv output |
| VDW Attractive | This value is the attractive energy between two atoms on different residues separated by distance, d. In the Rosetta `score.sc` output file, this value is called `fa_atr`. | rosetta_fa_atr |
| VDW Repulsive | This value is the repulsive energy between two atoms on different residues separated by distance, d. In the | rosetta_fa_rep |

| | | |
|---|---|---|
| | Rosetta `score.sc` output file, this value is called `fa_rep`. | |
| VDW Repulsive Intra Residue | This value is the repulsive energy between two atoms on the same residue separated by distance, d. In the Rosetta `score.sc` output file, this value is called `fa_intra_rep`. | rosetta_fa_intra_rep |
| Electrostatics | This value is the energy of interaction between two non-bonded charged atoms separated by distance, d. In the Rosetta `score.sc` output file, this value is called `fa_elec`. | rosetta_fa_elec |
| Solvation Isotropic | This value is the Gaussian exclusion implicit solvation energy between protein atoms in different residues. In the Rosetta `score.sc` output file, this value is called `fa_sol`. | rosetta_fa_sol |
| Solvation Anisotropic Polar Atoms | This value is the orientation-dependent solvation of polar atoms assuming ideal water geometry. In the Rosetta `score.sc` output file, this value is called `lk_ball_wtd`. | rosetta_lk_ball_wtd |
| Solvation Isotropic Intra Residue | This value is the Gaussian exclusion implicit solvation energy between protein atoms in the same residue. In the Rosetta `score.sc` output file, this value is called `fa_sol_intraR`. | rosetta_fa_intra_sol_xover4 |
| HB Long Range Backbone | This value is the energy of long range hydrogen bonds. In the Rosetta `score.sc` output file, this value is called `hbond_lr_bb`. | rosetta_hbond_lr_bb |
| HB Short Range Backbone | This value is the energy of short range hydrogen bonds. In the Rosetta `score.sc` output file, this value is called `hbond_sr_bb`. | rosetta_hbond_sr_bb |
| HB Backbone Sidechain | This value is the energy of backbone-side chain hydrogen bonds. In the Rosetta `score.sc` output file, this value is called `hbond_bb_sc`. | rosetta_hbond_bb_sc |
| HB Sidechain Sidechain | This value is the energy of side chain-side chain hydrogen bonds. In the Rosetta `score.sc` output file, this value is called `hbond_sc`. | rosetta_hbond_sc |
| Disulfide Bridges | This value is the energy of disulfide bridges. In the Rosetta `score.sc` output file, this value is called `dslf_fa13`. | rosetta_dslf_fa13 |
| Backbone Torsion Preference | This value is the probability of backbone ϕ, ψ angles given the amino acid type. In the Rosetta `score.sc` output file, this value is called `rama_prepro`. | rosetta_rama_prepro |
| Amino Acid Propensity | This value is the probability of amino acid identity given the backbone ϕ, ψ angles. In the Rosetta `score.sc` output file, this value is called `p_aa_pp`. | rosetta_p_aa_pp |
| Dunbrack Rotamer | This value is the probability that a chosen rotamer is native-like given backbone ϕ, ψ angles. In the Rosetta `score.sc` output file, this value is called `fa_dun`. | rosetta_fa_dun |
| Omega Penalty | This value is a backbone-dependent penalty for cis ω dihedrals that deviate from 0° and trans ω dihedrals that deviate from 180°. In the Rosetta `score.sc` output file, this value is called `omega`. | rosetta_omega |
| Open Proline Penalty | This value is a penalty for an open proline ring and proline ω bonding energy. In the Rosetta `score.sc` output file, this value is called `pro_close`. | rosetta_pro_close |
| Tyrosine χ3 Dihedral Angle Penalty | This value is a sinusoidal penalty for non-planar tyrosine χ3 dihedral angle. In the Rosetta `score.sc` output file, this value is called `yhh_planarity`. | rosetta_yhh_planarity |

*Table S7: This table provides a description of the metrics from the Rosetta programme and their variable names in the CSV output file.*

| **Metric Name** | **Metric Description** | **Variable Name in CSV Output** |
|---|---|---|
| ALA - Composition | The proportion of residues that are Alanine (ALA) in the structure. | composition_ALA |
| ARG - Composition | The proportion of residues that are Arginine (ARG) in the structure. | composition_ARG |
| ASN - Composition | The proportion of residues that are Asparagine (ASN) in the structure. | composition_ASN |
| ASP - Composition | The proportion of residues that are Aspartic Acid (ASP) in the structure. | composition_ASP |
| CYS - Composition | The proportion of residues that are Cysteine (CYS) in the structure. | composition_CYS |
| GLN - Composition | The proportion of residues that are Glutamine (GLN) in the structure. | composition_GLN |
| GLU - Composition | The proportion of residues that are Glutamic Acid (GLU) in the structure. | composition_GLU |
| GLY - Composition | The proportion of residues that are Glycine (GLY) in the structure. | composition_GLY |
| HIS - Composition | The proportion of residues that are Histidine (HIS) in the structure. | composition_HIS |
| ILE - Composition | The proportion of residues that are Isoleucine (ILE) in the structure. | composition_ILE |
| LEU - Composition | The proportion of residues that are Leucine (LEU) in the structure. | composition_LEU |
| LYS - Composition | The proportion of residues that are Lysine (LYS) in the structure. | composition_LYS |
| MET - Composition | The proportion of residues that are Methionine (MET) in the structure. | composition_MET |
| PHE - Composition | The proportion of residues that are Phenylalanine (PHE) in the structure. | composition_PHE |
| PRO - Composition | The proportion of residues that are Proline (PRO) in the structure. | composition_PRO |
| SER - Composition | The proportion of residues that are Serine (SER) in the structure. | composition_SER |
| THR - Composition | The proportion of residues that are Threonine (THR) in the structure. | composition_THR |
| TRP - Composition | The proportion of residues that are Tryptophan (TRP) in the structure. | composition_TRP |
| TYR - Composition | The proportion of residues that are Tyrosine (TYR) in the structure. | composition_TYR |
| VAL - Composition | The proportion of residues that are Valine (VAL) in the structure. | composition_VAL |
| UNK - Composition | The proportion of residues that are Unknown (UNK) in the structure. | composition_UNK |
| Number of Residues | The number of amino acid residues in the structure. | num_residues |
| Mass (Da) | The mass of the structure in daltons (Da). | mass |

*Table S8: This table provides a description of other metrics included in DE-STRESS and their variable names in the CSV output file.*

### Physicochemical properties can be used to predict protein production levels

| Top Contributors to PC1 | Top Contributors to PC2 |
|---|---|
| aggrescan3d_min_value | composition_GLY |
| composition_LYS | composition_GLN |
| composition_GLY | composition_VAL |
| composition_ASP | composition_TYR |
| composition_PRO | composition_PRO |
| composition_GLN | rosetta_hbond_bb_sc |
| aggrescan3d_avg_value | composition_THR |
| composition_THR | composition_LYS |
| composition_GLU | hydrophobic_fitness |
| rosetta_hbond_bb_sc | composition_HIS |

Table S9: Top 10 contributors to PC1 and PC2 for PCA space in figure 2B. For this PCA space the robust scaling method was used, amino acid composition metrics were included, and the mutual information score was used to select features.

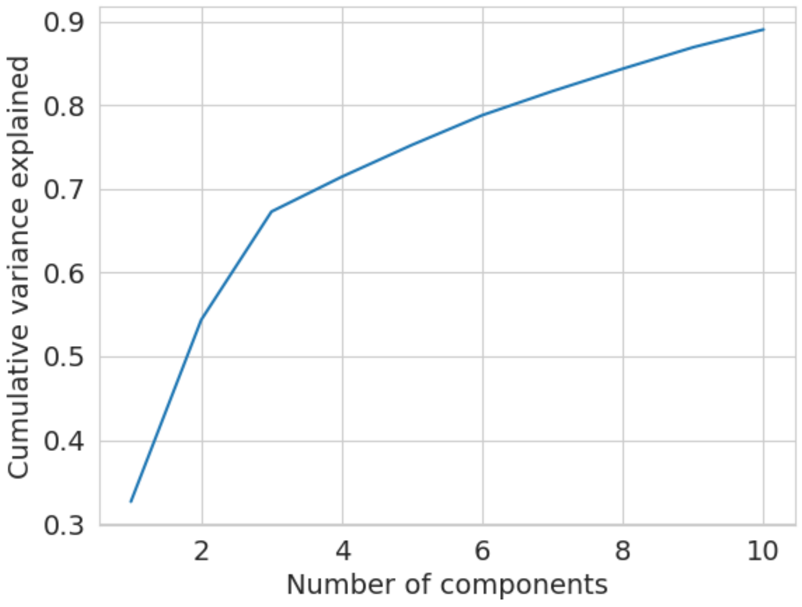

Figure S1: Cumulative variance explained by number of principal components for PCA space in figure 2B. For this PCA space the robust scaling method was used, amino acid composition metrics were included, and the mutual information score was used to select features.

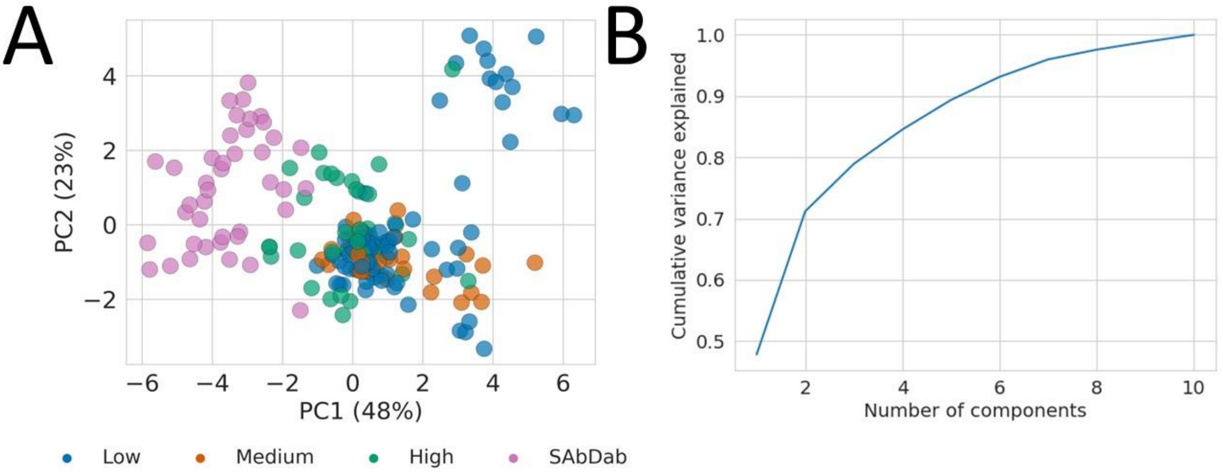

C

| Top Contributors to PC1 | Top Contributors to PC2 |
|---|---|
| rosetta_rama_prepro | aggrescan3d_min_value |
| aggrescan3d_min_value | rosetta_rama_prepro |
| rosetta_omega | aggrescan3d_avg_value |
| rosetta_hbond_bb_sc | rosetta_fa_sol |
| aggrescan3d_avg_value | rosetta_omega |

Figure S2: A) A plot of PC1 and PC2 for the scFv designs and 41 experimentally determined scFvs, along with the variance explained. For this PCA space the robust scaling method was used, amino acid composition metrics were excluded, and the mutual information score was used to select features. B) Cumulative variance explained by number of principal components. C) Table of the top contributors for PC1 and PC2.

| Scaling Method | Amino Acid Composition Metrics | Feature Selection Method | Mean Cross Validation ROC AUC | Mean Cross Validation Precision | Mean Cross Validation Recall |
|---|---|---|---|---|---|
| Minmax | Included | Random Forest | 0.762 | 0.635 | 0.545 |
| Standard | Included | Random Forest | 0.758 | 0.628 | 0.535 |
| Robust | Included | Random Forest | 0.757 | 0.625 | 0.538 |
| Standard | Excluded | Random | 0.750 | 0.624 | 0.536 |

| | | | | | |
|---|---|---|---|---|---|
| | | Forest | | | |
| Minmax | Excluded | Random Forest | 0.746 | 0.623 | 0.532 |
| Robust | Excluded | Random Forest | 0.744 | 0.626 | 0.527 |
| Robust | Included | Mutual Information | 0.743 | 0.647 | 0.518 |
| Minmax | Excluded | Mutual Information | 0.742 | 0.612 | 0.539 |
| Standard | Excluded | Mutual Information | 0.741 | 0.604 | 0.536 |
| Robust | Excluded | Mutual Information | 0.740 | 0.610 | 0.542 |
| Standard | Included | Mutual Information | 0.735 | 0.646 | 0.525 |
| Minmax | Included | Mutual Information | 0.734 | 0.643 | 0.523 |

Table S10: Averaged validation metrics for the naive bayes models fitted using 10 repetitions of 5-fold validation on different training sets. These metrics are sorted by the mean ROC AUC score.

| Scaling Method | Amino Acid Composition Metrics | Feature Selection Method | Test ROC AUC | Test Precision | Test Recall |
|---|---|---|---|---|---|
| Standard | Included | Random Forest | 0.785 | 0.683 | 0.604 |
| Robust | Included | Random Forest | 0.785 | 0.683 | 0.604 |
| Minmax | Included | Random Forest | 0.785 | 0.683 | 0.604 |
| Standard | Included | Mutual Information | 0.765 | 0.728 | 0.604 |
| Robust | Included | Mutual Information | 0.765 | 0.728 | 0.604 |
| Minmax | Included | Mutual Information | 0.765 | 0.728 | 0.604 |
| Standard | Excluded | Mutual Information | 0.757 | 0.562 | 0.521 |
| Robust | Excluded | Mutual Information | 0.757 | 0.562 | 0.521 |
| Minmax | Excluded | Mutual Information | 0.757 | 0.562 | 0.521 |
| Standard | Excluded | Random Forest | 0.733 | 0.531 | 0.500 |
| Robust | Excluded | Random Forest | 0.733 | 0.531 | 0.500 |

| | | | | | |
|---|---|---|---|---|---|
| Minmax | Excluded | Random Forest | 0.733 | 0.531 | 0.500 |

Table S11: Validation metrics for the naive bayes models on the different test sets. These metrics are sorted by the mean ROC AUC score.

| **Including amino acid composition** | **Excluding amino acid composition** |
|---|---|
| composition_GLN | aggrescan3d_max_value |
| hydrophobic_fitness | rosetta_fa_sol |
| composition_VAL | aggrescan3d_avg_value |
| composition_ASP | rosetta_fa_atr |
| composition_GLU | rosetta_hbond_bb_sc |
| composition_PRO | aggrescan3d_min_value |
| composition_GLY | evoef2_total |
| aggrescan3d_max_value | rosetta_fa_intra_sol_xover |
| composition_LEU | rosetta_fa_elec |
| aggrescan3d_avg_value | rosetta_fa_dun |
| composition_HIS | |
| rosetta_fa_intra_sol_xover4 | |
| rosetta_fa_sol | |
| composition_THR | |

Table S12: Random Forest selected features including and excluding amino acid composition metrics. The same features were found for the minmax, standard and robust scaling methods. Similar features were also found by using the mutual information score, which is shown in table S13.

| **Including amino acid composition** | **Excluding amino acid composition** |
|---|---|
| composition_HIS | aggrescan3d_max_value |
| composition_GLN | evoef2_total |
| composition_GLU | aggrescan3d_min_value |
| hydrophobic_fitness | aggrescan3d_avg_value |
| composition_PRO | rosetta_hbond_bb_sc |
| composition_LYS | rosetta_omega |
| composition_VAL | rosetta_fa_atr |
| composition_ASP | rosetta_rama_prepro |
| aggrescan3d_max_value | rosetta_p_aa_pp |
| composition_GLY | rosetta_fa_sol |
| aggrescan3d_min_value | |
| composition_PHE | |
| aggrescan3d_avg_value | |

| |
|---|
| composition_THR |
| composition_TYR |
| rosetta_hbond_bb_sc |
| composition_ILE |
| composition_TRP |
| composition_LEU |

Table S13: Mutual information selected features including and excluding amino acid composition metrics. The same features were found for the different scaling methods.

#### Large-scale analysis of physicochemical properties across half a million predicted protein structures

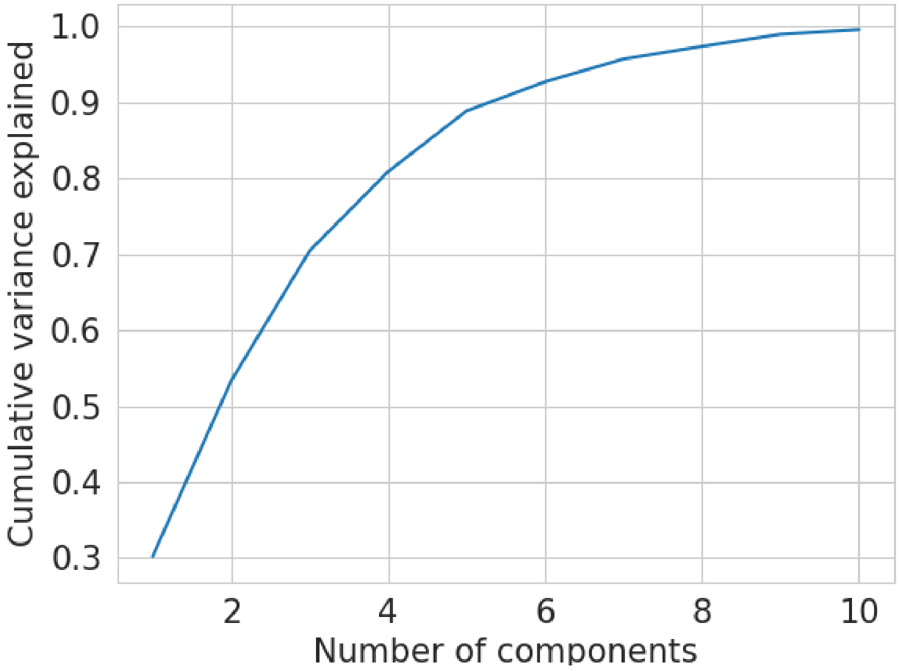

Figure S3: Cumulative variance explained by number of principal components for the AF2 structural model PCA space in figure 3A. For this PCA space the minmax scaling method was used and amino acid composition metrics were excluded.

#### AF2 structural models – Min max scaling

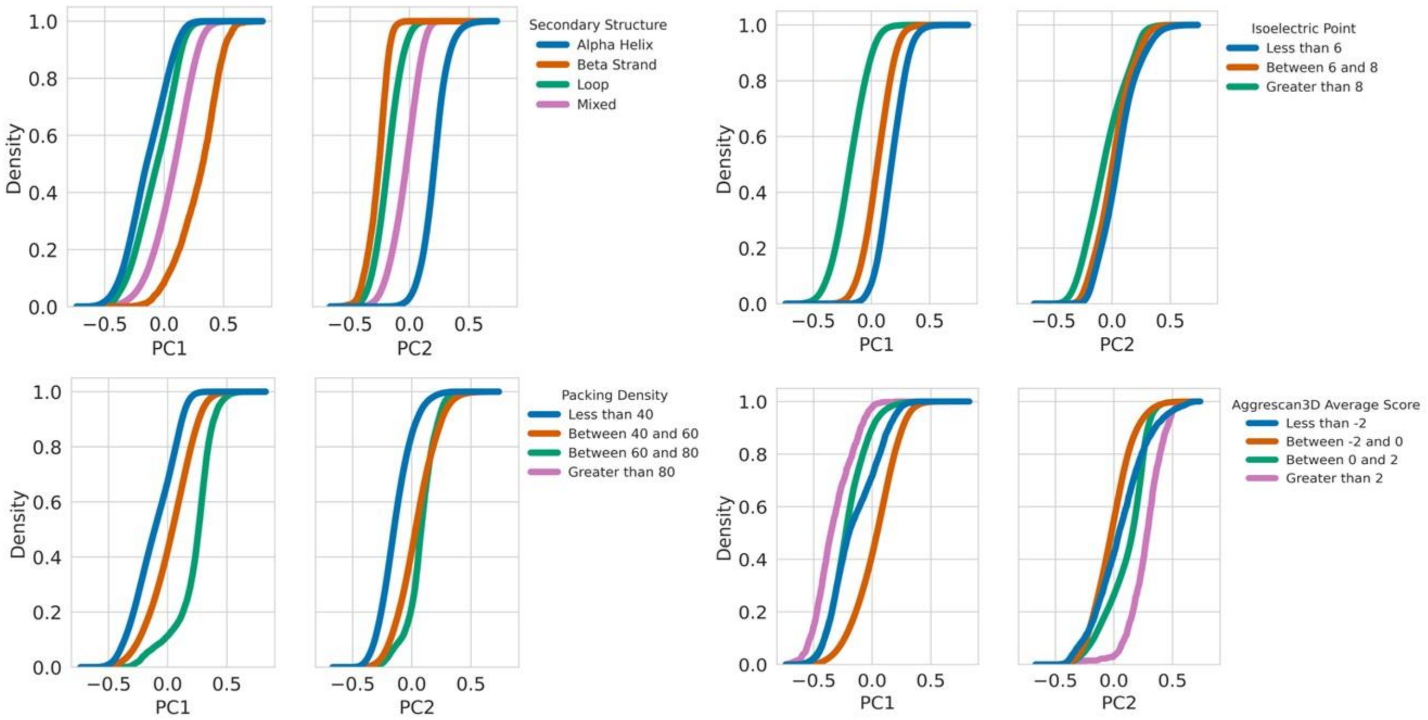

Figure S4: Cumulative histograms of different metrics, across PC1 and PC2, for the physicochemical properties of 564,442 AF2 structural models in figure 3A. The minmax scaling method was used for this PCA space.

#### AF2 structural models – Robust scaling

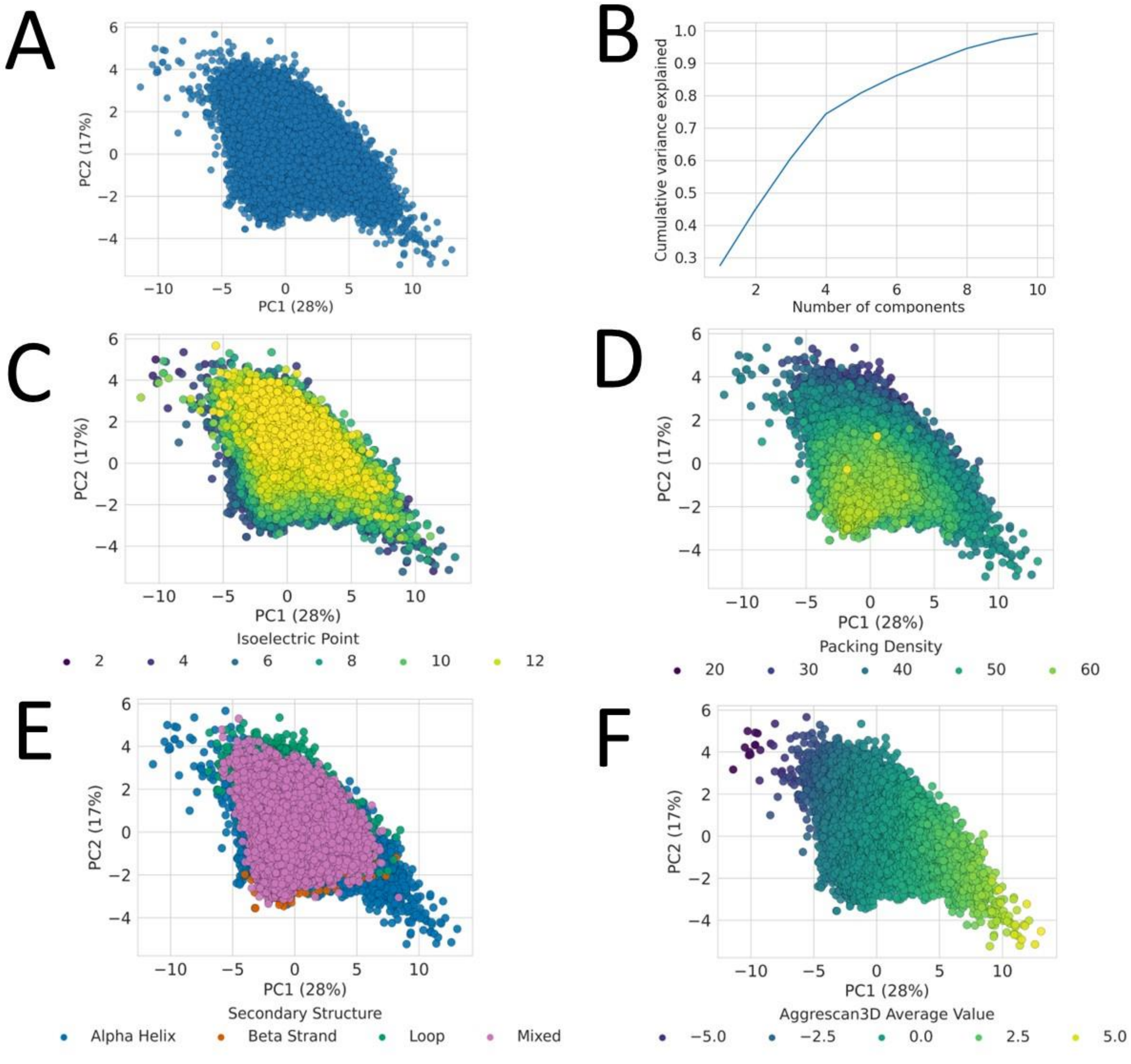

Figure S5: A), C), D), E), F) shows PC1 and PC2 for the physicochemical properties of the 564,442 AF2 structural models and how different metrics vary across this space, while B) shows the cumulative variance explained by number of components. For this space the robust scaling method was used, and amino acid composition metrics were excluded.

| Top Contributors to PC1 | Top Contributors to PC2 |
|---|---|
| aggrescan3d_avg_value | rosetta_fa_dun |
| aggrescan3d_min_value | rosetta_hbond_sc |
| rosetta_fa_intra_sol_xover4 | aggrescan3d_min_value |
| aggrescan3d_max_value | rosetta_fa_intra_sol_xover4 |
| budeff_charge | rosetta_hbond_lr_bb |

Table S14: Top 5 contributors to PC1 and PC2 for AF2 structural model PCA space in figure S4. For this PCA space the robust scaling method was used, and amino acid composition metrics were excluded.

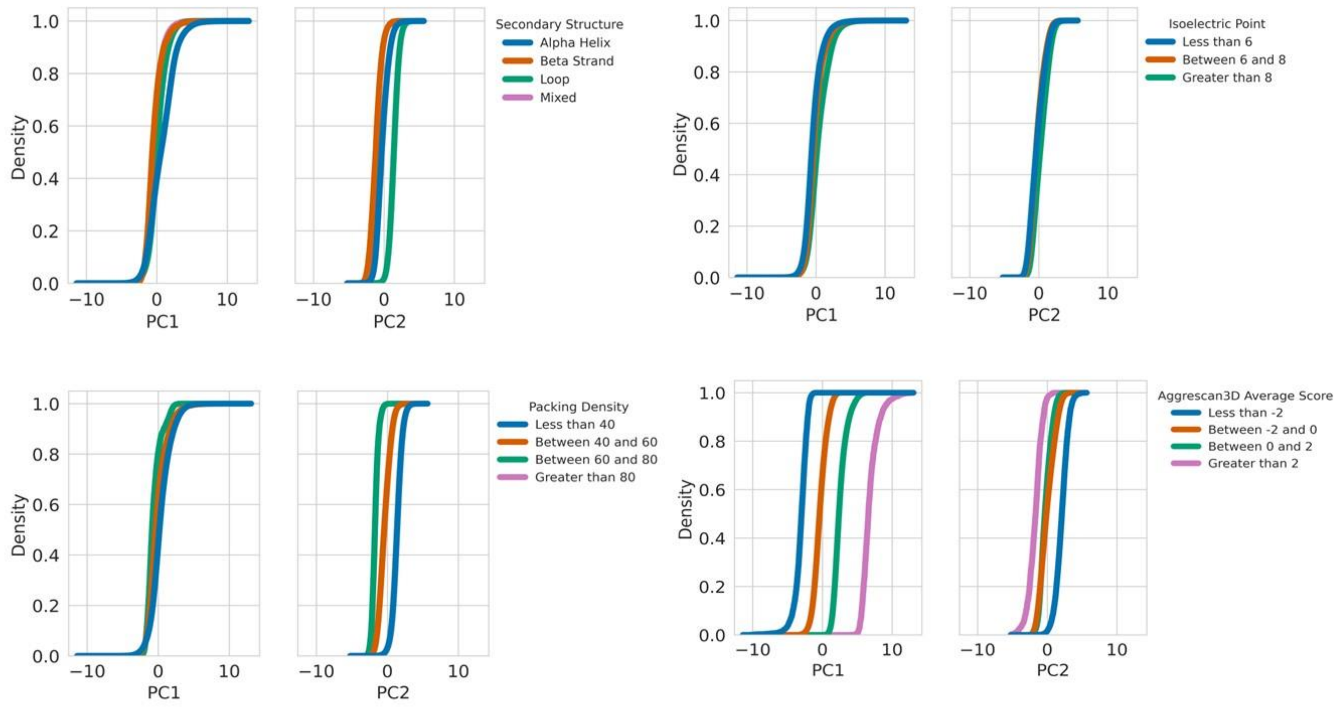

Figure S6: Cumulative histograms of different metrics, across PC1 and PC2, for the physicochemical properties of 564,442 AF2 structural models in figure S4. The robust scaling method was used for this PCA space.

#### PDB structures – Min max scaling

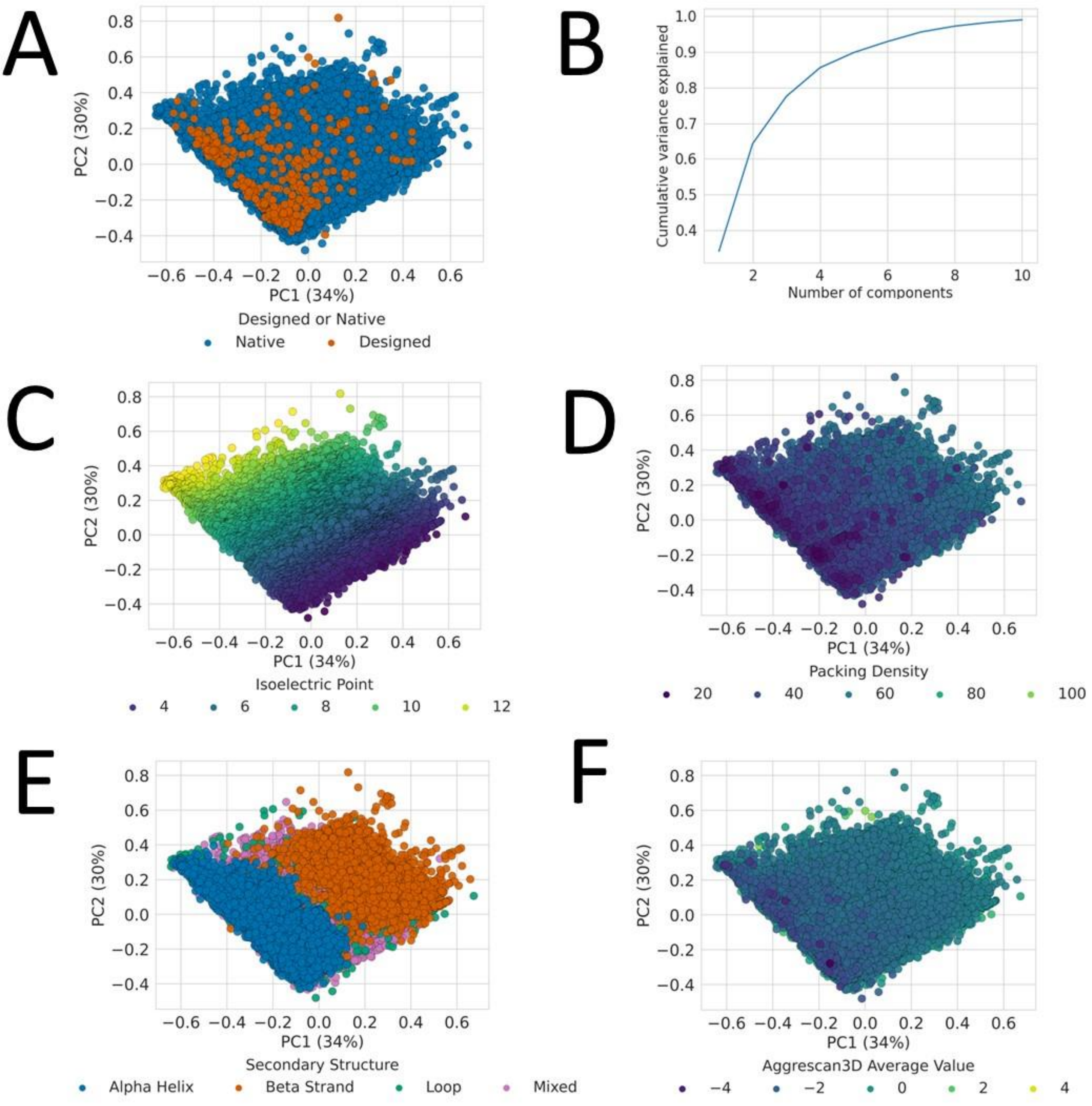

Figure S7: A), C), D), E), F) shows PC1 and PC2 for the physicochemical properties of the PDB and how different metrics vary across this space, while B) shows the cumulative variance explained by number of components. For this space the minmax scaling method was used, and amino acid composition metrics were excluded.

| **Top Contributors to PC1** | **Top Contributors to PC2** |
|---|---|
| rosetta_hbond_lr_bb | isoelectric_point |
| isoelectric_point | rosetta_hbond_lr_bb |
| rosetta_hbond_bb_sc | rosetta_hbond_bb_sc |
| rosetta_hbond_sc | rosetta_fa_sol |
| packing_density | rosetta_lk_ball_wtd |

Table S15: Top 5 contributors to PC1 and PC2 for PDB PCA space in figure S6. For this PCA space the minmax scaling method was used, and amino acid composition metrics were excluded.

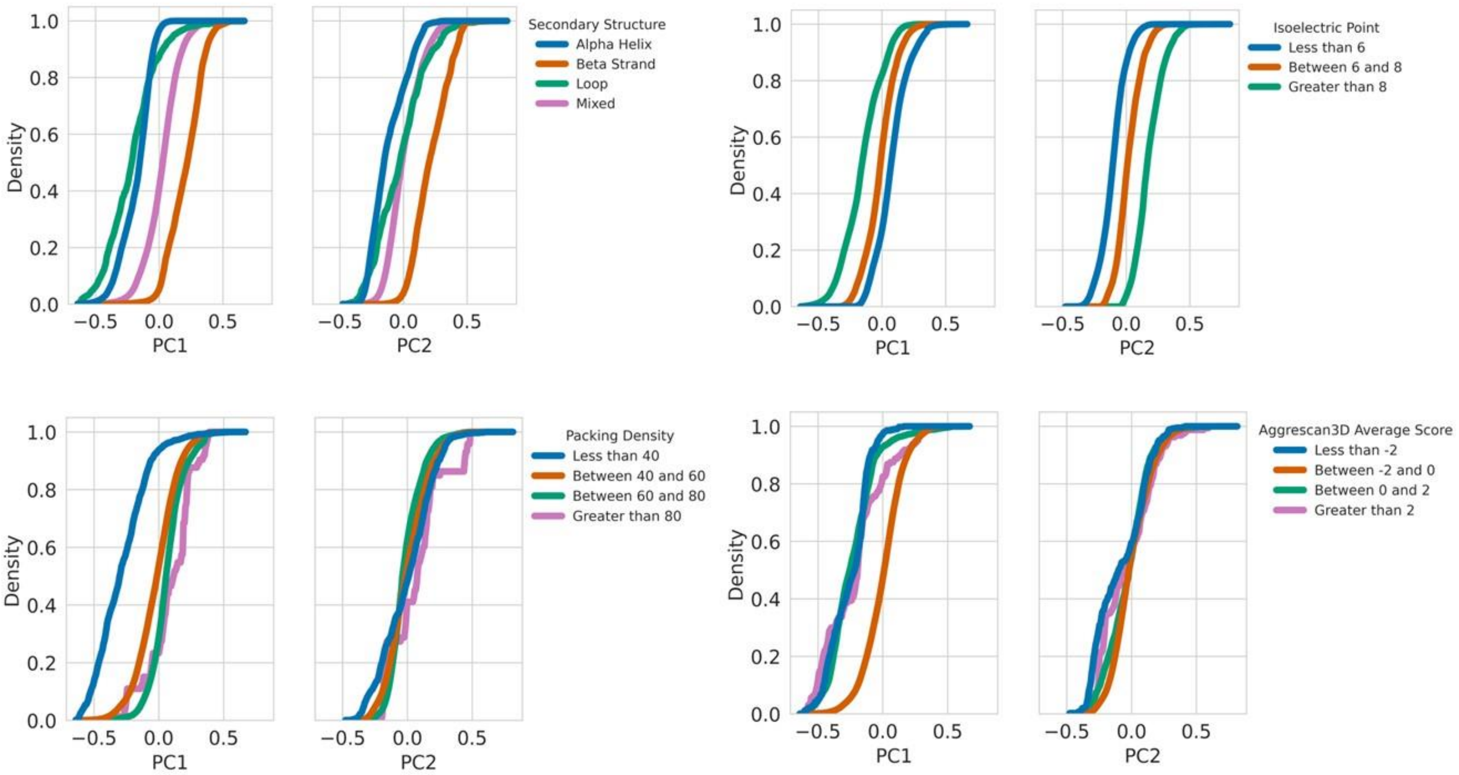

Figure S8: Cumulative histograms of different metrics, across PC1 and PC2, for the physicochemical properties of PDB structures in figure S6. The minmax scaling method was used for this PCA space.

#### PDB structures – Robust scaling

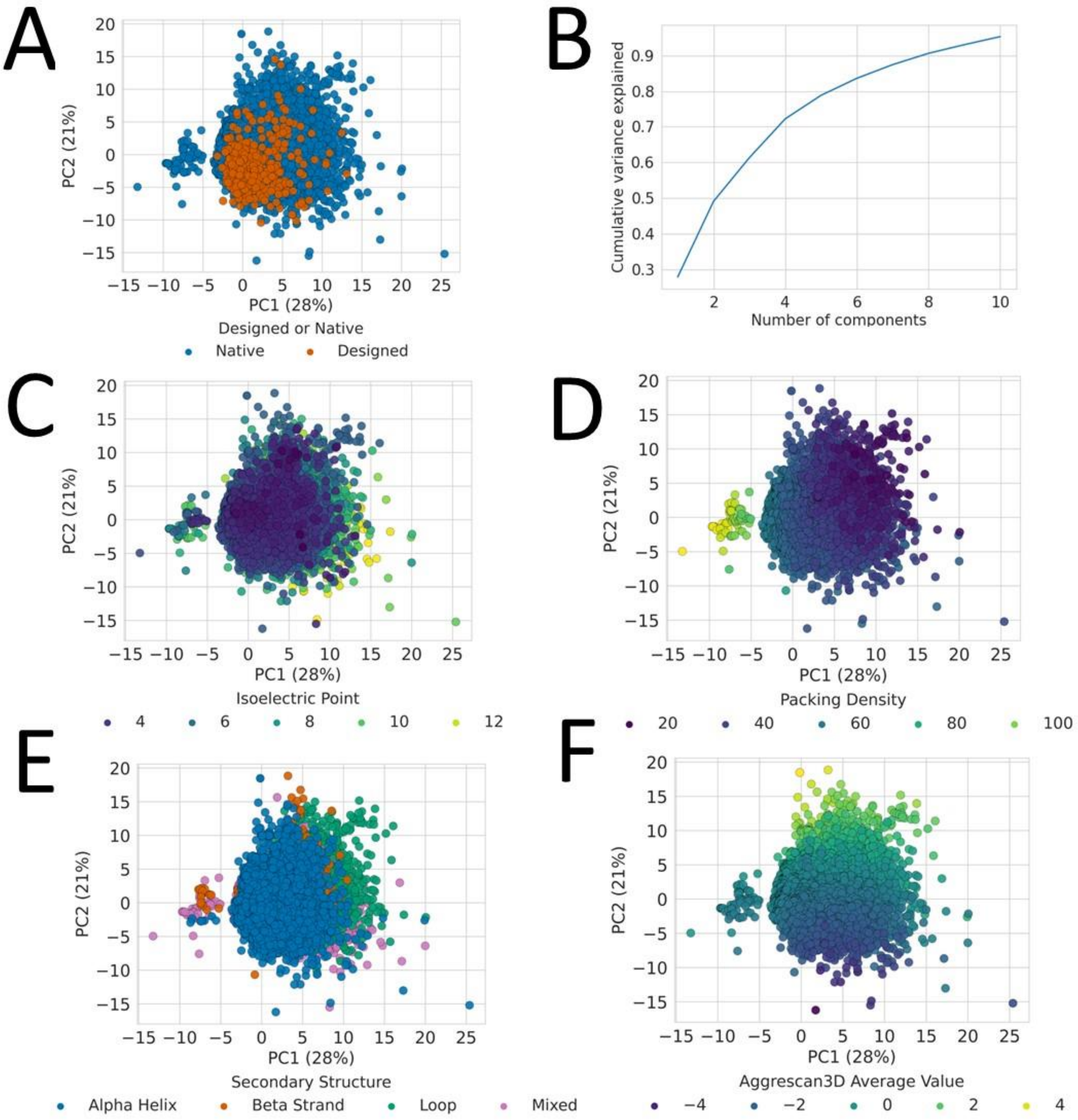

Figure S9: A), C), D), E), F) shows PC1 and PC2 for the physicochemical properties of the PDB and how different metrics vary across this space, while B) shows the cumulative variance explained by number of components. For this space the robust scaling method was used, and amino acid composition metrics were excluded.

| **Top Contributors to PC1** | **Top Contributors to PC2** |
|---|---|
| rosetta_fa_dun | aggrescan3d_avg_value |
| packing_density | rosetta_fa_intra_sol_xover4 |
| evoef2_intraR_total | aggrescan3d_min_value |
| rosetta_hbond_bb_sc | aggrescan3d_max_value |
| rosetta_hbond_sc | rosetta_fa_sol |

Table S16: Top 5 contributors to PC1 and PC2 for PDB PCA space in figure S7. For this PCA space the robust scaling method was used, and amino acid composition metrics were excluded.

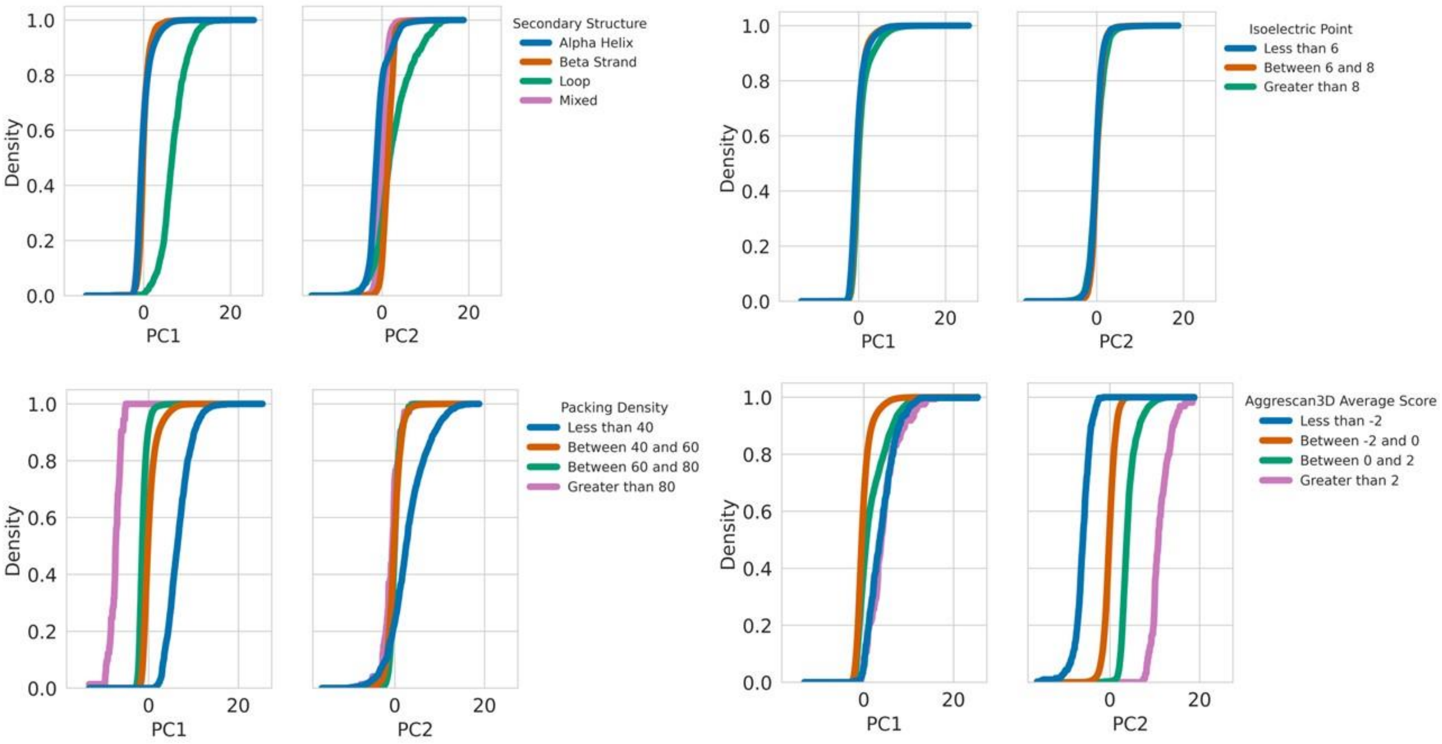

Figure S10: Cumulative histograms of different metrics, across PC1 and PC2, for the physicochemical properties of PDB structures in figure S7. The robust scaling method was used for this PCA space.

### Physicochemical properties distinguish eukaryotic and prokaryotic organisms

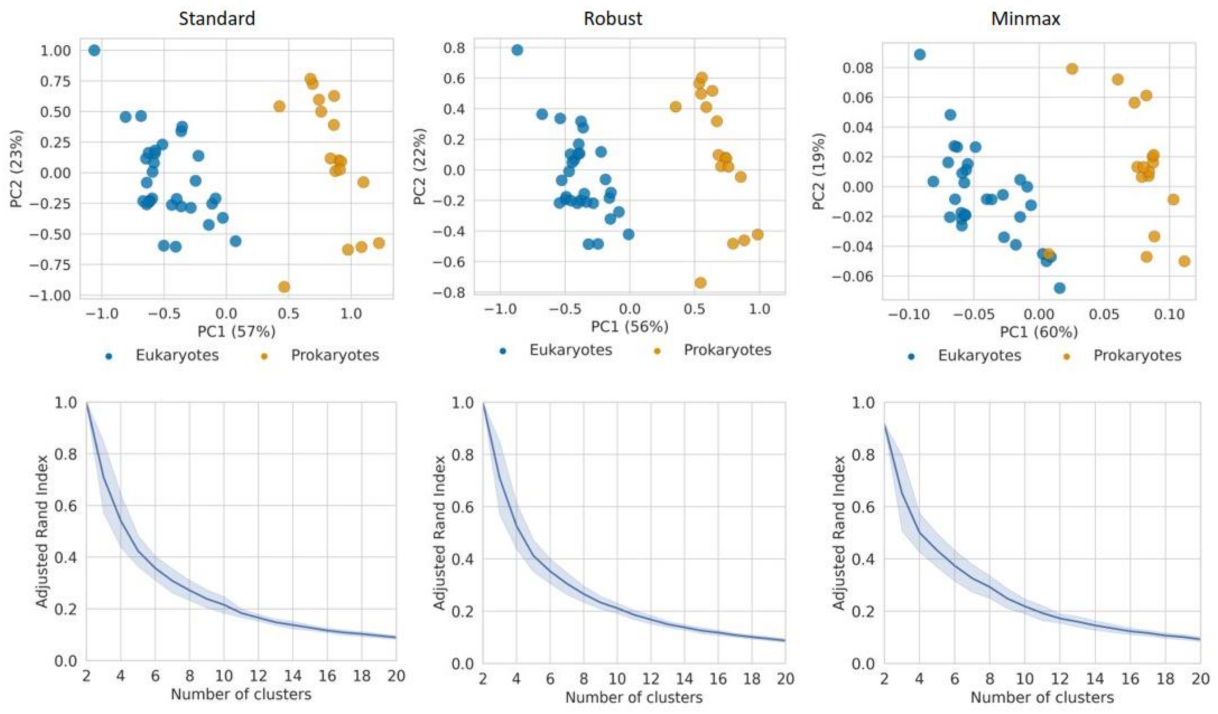

Figure S11: The average properties for each organism across PC1 and PC2, along with the variance explained for different scaling methods. Also, the mean and standard deviation of the adjusted rand index against the eukaryote and prokaryote groups, for 100 random initialisations of K-means, and different numbers of clusters.

| **Top Contributors to PC1** | **Top Contributors to PC2** |
|---|---|
| rosetta_fa_dun | rosetta_fa_intra_sol_xover4 |
| rosetta_fa_elec | rosetta_fa_elec |
| aggrescan3d_min_value | rosetta_lk_ball_wtd |
| aggrescan3d_max_value | rosetta_fa_dun |
| rosetta_hbond_bb_sc | rosetta_hbond_sr_bb |

Table S17: Top 5 contributors to PC1 and PC2 of the average properties for each organism using the standard scaling method.

| Top Contributors to PC1 | Top Contributors to PC2 |
|---|---|
| rosetta_fa_dun | rosetta_fa_intra_sol_xover4 |
| aggrescan3d_min_value | rosetta_fa_elec |
| rosetta_fa_elec | rosetta_lk_ball_wtd |
| aggrescan3d_max_value | rosetta_hbond_sr_bb |
| budeff_charge | rosetta_fa_dun |

Table S18: Top 5 contributors to PC1 and PC2 of the average properties for each organism using the robust scaling method.

| Top Contributors to PC1 | Top Contributors to PC2 |
|---|---|
| rosetta_fa_elec | rosetta_fa_elec |
| rosetta_fa_dun | isoelectric_point |
| aggrescan3d_max_value | rosetta_fa_intra_sol_xover4 |
| rosetta_hbond_sc | rosetta_hbond_sr_bb |
| rosetta_hbond_bb_sc | rosetta_lk_ball_wtd |

Table S19: Top 5 contributors to PC1 and PC2 of the average properties for each organism using the minmax scaling method.

#### Reconstructing the tree of life from model-derived properties

| Scaling Method | Linkage Method | Clustering information distance to NCBI Taxonomic Tree |
|---|---|---|
| Minmax | Average | 0.43 |
| Standard | Average | 0.43 |
| Minmax | Complete | 0.43 |
| Minmax | Ward | 0.44 |
| Standard | Ward | 0.46 |
| Robust | Ward | 0.47 |
| Robust | Average | 0.47 |
| Robust | Complete | 0.47 |
| Standard | Complete | 0.48 |
| Minmax | Single | 0.49 |
| Standard | Single | 0.53 |
| Robust | Single | 0.53 |

Table S20: Clustering information distance against NCBI Taxonomic tree for different scaling and linkage metrics.

| Organism name | Volume of structures | Volume of non-redundant structures | Volume of high-quality models (Avg PLDDT >= 70%) |
|---|---|---|---|
| Ajellomyces capsulatus | 9,199 | 8,666 | 4,575 |
| Arabidopsis thaliana | 27,427 | 14,455 | 8,503 |
| Brugia malayi | 8,731 | 6,876 | 4,311 |
| Caenorhabditis elegans | 19,671 | 13,948 | 8,655 |
| Campylobacter jejuni | 1,620 | 1,536 | 1,499 |
| Candida albicans | 5,974 | 5,325 | 3,770 |
| Cladophialophora carrionii | 11,170 | 9,825 | 6,192 |
| Danio rerio | 24,656 | 13,802 | 9,382 |
| Dictyostelium discoideum | 12,622 | 10,129 | 5,396 |
| Dracunculus medinensis | 10,834 | 9,864 | 6,391 |
| Drosophila melanogaster | 13,455 | 10,853 | 6,520 |
| Enterococcus faecium | 2,627 | 2,264 | 2,152 |
| Escherichia coli | 4,363 | 3,657 | 3,515 |
| Fonsecaea pedrosoi | 12,509 | 9,768 | 6,589 |

| | | | |
|---|---|---|---|
| Glycine max | 55,798 | 23,349 | 12,422 |
| Haemophilus influenzae | 1,661 | 1,583 | 1,545 |
| Helicobacter pylori | 1,538 | 1,460 | 1,352 |
| Homo sapiens | 23,327 | 13,755 | 8,489 |
| Klebsiella pneumoniae | 5,727 | 4,782 | 4,409 |
| Leishmania infantum | 7,914 | 7,336 | 3,941 |
| Madurella mycetomatis | 9,561 | 8,325 | 5,809 |
| Methanocaldococcus jannaschii | 1,773 | 1,633 | 1,545 |
| Mus musculus | 21,562 | 12,577 | 8,090 |
| Mycobacterium leprae | 1,602 | 1,520 | 1,344 |
| Mycobacterium tuberculosis | 3,988 | 3,336 | 3,043 |
| Mycobacterium ulcerans | 9,033 | 7,876 | 6,000 |
| Neisseria gonorrhoeae | 2,106 | 1,967 | 1,713 |
| Nocardia brasiliensis | 8,372 | 6,087 | 5,685 |
| Onchocerca volvulus | 1,459 | 1,391 | 926 |
| Oryza sativa | 41,894 | 30,164 | 10,980 |

| | | | |
|---|---|---|---|
| Paracoccidioides lutzii | 8,794 | 8,358 | 4,532 |
| Plasmodium falciparum | 5,115 | 4,597 | 2,053 |
| Pseudomonas aeruginosa | 5,556 | 4,366 | 4,207 |
| Rattus norvegicus | 19,258 | 12,220 | 7,874 |
| Saccharomyces cerevisiae | 6,038 | 5,116 | 3,584 |
| Salmonella typhimurium | 4,526 | 3,834 | 3,660 |
| Schistosoma mansoni | 13,856 | 10,505 | 5,401 |
| Schizosaccharomyces pombe | 5,112 | 4,539 | 3,429 |
| Shigella dysenteriae | 3,893 | 3,189 | 3,003 |
| Sporothrix schenckii | 8,652 | 7,691 | 4,801 |
| Staphylococcus aureus | 2,888 | 2,570 | 2,382 |
| Streptococcus pneumoniae | 2,030 | 1,834 | 1,751 |
| Strongyloides stercoralis | 12,609 | 10,233 | 6,243 |
| Trichuris trichiura | 9,563 | 8,581 | 5,662 |
| Trypanosoma brucei | 8,491 | 7,592 | 4,354 |
| Trypanosoma cruzi | 19,036 | 10,280 | 6,004 |

| | | | |
|---|---|---|---|
| Wuchereria bancrofti | 12,721 | 12,315 | 7,245 |
| Zea mays | 38,914 | 21,881 | 10,206 |
| **Total** | **549,225** | **387,810** | **241,134** |

Table S21: Table of the 48 different organisms, along with initial volume of protein structures downloaded from AlphaFold DB and then the volumes after using MMSeq2 to remove redundant proteins, and after removing low quality AF2 structural models.
